# Attentive-SPIDNA: Attention-based neural networks for population genetics

**DOI:** 10.64898/2026.04.15.718687

**Authors:** Théophile Sanchez, Pierre Jobic, Cyril Regan, Paul Verdu, Guillaume Charpiat, Flora Jay

**Author notes:** These authors contributed equally (Théophile Sanchez, Pierre Jobic).

## Abstract

Artificial neural networks (ANNs) have recently offered new perspectives to solve inference problems from high dimensional data in numerous scientific fields, but it is yet unclear which architectures are the most suited to genomic data. Here, we present a new ANN architecture integrating attention mechanisms to infer effective population size history from genomic data. Built upon our previous exchangeable architecture SPIDNA, Attentive-SPIDNA adds attention layers that allow computing more expressive and complex features from combinations of haplotypes. The contribution of each haplotype to the features is learned automatically and depends on its content and affinity with the other haplotypes. Likewise, we use this mechanism to automatically perform a voting scheme that aggregates predictions from different genomic regions. This new architecture outperforms approximate Bayesian computation and previously published neural networks while relying directly on raw genetic data and being invariant to haplotype permutation in the input. As a proof-of-concept, we use this architecture to infer the effective population size history of 54 populations from the HGDP dataset (Bergström et al, 2020). This application highlights the ability of the network to handle data with a varying number of haplotypes and to quickly perform predictions for datasets including numerous populations. Therefore, the proposed mechanism could be integrated to various neural networks solving population genetics tasks.

## Introduction

Inferring the evolutionary past from genomic data is a long-standing question in the population genetics field, and a variety of approaches have been proposed until now. From the initial model-based likelihood inference likelihood-free inference approaches based on simulation, all have in common to leverage the information imprinted into complex genomic patterns of multiple individuals (or at least one diploid individual). A notable development in the past decade is the focus on how exactly the information can be retrieved. We have observed a transition from handcrafted feature design, commonly called summary statistics, to semi-automatic or automatic feature construction.

This shift began with efforts to enhance commonly used features by combining them using various linear (Chun and Keles, 2010; Wegmann et al, 2009) and non-linear methods. These operations can be performed in numerous way, leading to the natural adoption of supervised machine learning when the number of summary statistics inherently increased with the advent of genome-scale data. Support Vector Machines (SVM) or Random Forests (RF) and their extensions have been and continue to be popular choices for this purpose (e.g. Schrider and Kern, 2016; Smith et al, 2017; Raynal et al, 2019) (see Schrider and Kern, 2018, for a review). In recent years, some studies replaced classical machine learning with deep learning. The goal remains to learn complex interactions between summary statistics to optimize information extraction. In this context, deep learning has been used for direct prediction (Sheehan and Song, 2016; Kern and Schrider, 2018; Hejase et al, 2022; Villanea and Schraiber, 2018; Quelin et al, 2025) or combined with Approximate Bayesian Computation (ABC) schemes (Mondal et al, 2019; Sanchez et al, 2021; Kirschner et al, 2022) as suggested by Jiang et al (2017).

Finally, end-to-end deep learning approaches have emerged, leveraging the hierarchical learning capabilities of deep neural networks to automatically extract features. These networks, in theory, can approximate any function, as demonstrated by Cybenko (1989) and Hornik et al (1989). For comprehensive information on deep learning approaches in evolutionary studies, we refer readers to recent reviews (Borowiec et al, 2022; Korfmann et al, 2023; Yelmen and Jay, 2023; Huang et al, 2024). In this work, we focus specifically on the issue of exchangeability. SNP matrices, which are frequently used as inputs for artificial neural networks in population genetics, possess an exchangeable (or permutation-invariant) property (Chan et al, 2018; Sanchez et al, 2021; Flagel et al, 2018; Torada et al, 2019). This means that the order of rows (representing haploid or diploid individuals) should not affect the prediction of population parameters such as effective population sizes, selection coefficients, and recombination rates, given that there is typically no inherent ordering in individual data collection. Three approaches have been explored for applying networks to raw population genetics data: (i) Non-exchangeable networks: these networks can become invariant only though training (Kern and Schrider, 2018; Hejase et al, 2022; Villanea and Schraiber, 2018); (ii) Ordering operations: some studies apply a systematic ordering operation to input matrices (Flagel et al, 2018; Torada et al, 2019); (iii) Invariance by design: architectures such as convolutional neural networks (CNNs) (Chan et al, 2018; Sanchez et al, 2021; Wang et al, 2021) or graph-based networks (Korfmann et al, 2024) can be constructed to be inherently invariant. Graph encoding has the advantage of inputting a lot of prior information regarding the input data structure with the drawback that, in reality, this graph is unknown and has to be learned (along the sequence). The exchangeable networks proposed by Chan et al (2018), Wang et al (2021) and Sanchez et al (2021) have in common the use of a simple invariant function, such as the sum or the mean, although other statistical moments could be used. Initially, these network aggregated individual-based equivariant features into invariant features only once at the end of the network (Zaheer et al, 2017; Chan et al, 2018; Wang et al, 2021). Following an original idea of Lucas et al (2018), who demonstrated its capability of approximating any invariant function, Sanchez et al (2021) introduced a novel architecture that repeatedly combines global invariant information with the local individual-based information at each block, facilitating the learning of complex invariant functions.

However, simply averaging over all individuals can create an information bottleneck. To address this issue, our study proposes a novel exchangeable architecture that incorporates attention mechanisms, enabling the automatic construction of expressive permutation-invariant features. Since the introduction of the self-attention mechanism in the transformer architecture (Vaswani et al, 2017), attention-based neural networks have shown remarkable capabilities in various domains. They have been highly successful in natural language processing tasks (Devlin et al, 2019; Brown et al, 2020), image sampling and reconstruction (Chen et al, 2020), image generation (Ramesh et al, 2021; Ho et al, 2020), protein structure prediction and relationships (Jumper et al, 2021; Rao et al, 2021) and other genome-based prediction tasks (e.g. Nesterenko et al, 2025) (see Consens et al (2023) and Choi and Lee (2023) for a review). Similar to recurrent layers, attention mechanisms are designed to process sequences, but they offer the advantage of parallelizing computation across all sequence elements rather than processing them sequentially. Unlike convolutional layers, attention mechanisms connect all sequence elements to each other, preventing the loss of information between distant elements and thereby handling long-range dependencies more effectively. Additionally, unlike fully connected layers, attention mechanisms weight the input data based on both their composition and context, resulting in more flexible functions that vary for each input. In our study, attention is applied to the transformed representations of either individuals or genomic regions, enabling a dynamic weighted average of their features.

In this work, we demonstrate that attention mechanisms can greatly improve performance in evolutionary inference by focusing on the complex task of reconstructing past effective population size histories from raw whole genome sequences. To illustrate this, we developed a new exchangeable architecture called Attentive-SPIDNA, that leverages convolution and attention mechanisms for both aggregating information within genomic regions and between genomic regions. This approach significantly improves performance on simulated datasets compared to a wide variety of methods including non-exchangeable and exchangeable networks, both with and without the use of summary statistics and ABC. Finally, we apply our novel approach to analyze the recently resequenced 54 worldwide populations of the HGDP dataset (Bergström et al, 2020). We emphasize that the concept behind the proposed mechanisms has a general implication, and could be incorporated into existing or future networks for evolutionary inference from pools of varying numbers of sequences.

## Results

### A more expressive exchangeable network for evolutionary inference

We developed a new architecture, Attentive-SPIDNA, to increase the expressivity of exchangeable networks (Chan et al, 2018; Sanchez et al, 2021) while retaining their invariance property. One bottleneck of these architectures stems from their equivariant component, which relies solely on a simple mean over the haplotype dimension of the data flowing through the network. The mean, however, is not a very rich descriptor of a distribution of features over individuals. We thus are interested in incorporating additional statistical descriptors that better capture information about the distribution of individuals. While Lucas et al (2018) demonstrates that the mean is sufficiently informative in the sense that all permutation-invariant functions can be expressed with a Deep Sets architecture based on the mean, provided there are enough layers, the number of required layers is unknown. One can hope that with richer statistical descriptors in each layer, the network can be more efficient and shallower.

To implement this idea, Attentive-SPIDNA integrates an attention mechanism called attention hub into the original SPIDNA (Sanchez et al, 2021). The intuition behind this mechanism, is that it allows the network to learn complex invariant features from subsets of individuals or from individuals contributing to these features to varying extent. It functions as a concatenation of multiple learned weighted means, with weights depending on the individuals’ features rather than their position in the SNP matrix. This subtlety renders the operation invariant to permutation of the individuals, but also directly applicable to any number of individuals without the need for retraining. Additionally, a second reversed attention mechanism is applied at each block to mix the invariant features back to individual equivariant information.

Secondly, we introduce a novel method for aggregating the predictions obtained from multiple genomic regions via a secondary a second attention hub mechanism. This approach enables the inference of demographic parameters from a collection of regions, regardless of their ordering, while using richer aggregations than a simple mean.

### Benchmarking ABC approaches, non-exchangeable and exchangeable neural networks

To determine how the Attentive-SPIDNA architecture ranks against the methods previously compared in Sanchez et al (2021), we used the same dataset of populations with a set of broad demographic priors and step-wise constant effective population sizes (21 steps), simulated using the scheme originally developed by Boitard et al (2016). We used the mean squared error (MSE) both as evaluation metric to compare methods and as loss function.

Previous methods encompass four ABC algorithms and one multilayer perceptron (MLP) based on summary statistics only, one MLP based on SNPs, variations of two non-exchangeable CNNs based on SNPs (one Custom CNN designed by Sanchez et al (2021), four “Flagel CNN” designed by Flagel et al (2018)), five variations of the exchangeable SPIDNA network based on SNPs and 16 combinations of SPIDNA with ABC (including summary statistics or not). We newly trained three versions of our attention-based architecture, Attentive-SPIDNA, on the same simulated “benchmark” dataset and compared training, validation and test errors for the 34 methods (See Table 1 and Table S1).

**Table 1:**
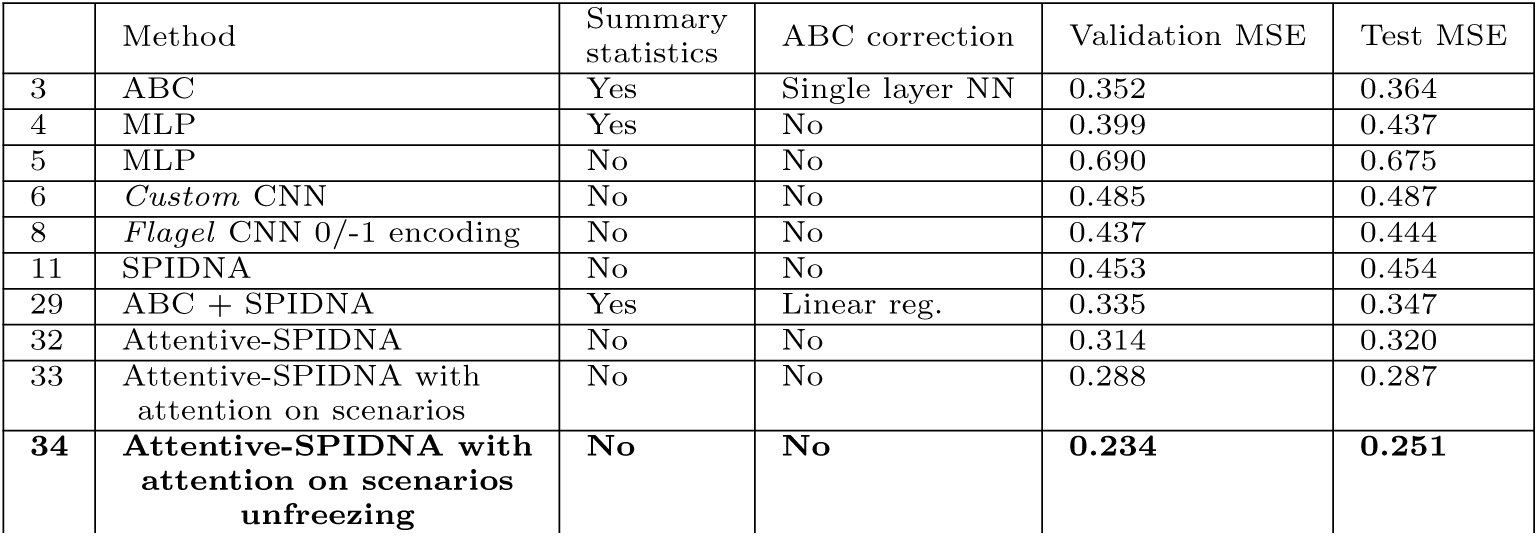
Summary of the prediction errors of each method on the simulated benchmark dataset. For the complete set of results, please refer to Table S1. All method’s hyperparameters, but Attentive-SPIDNA’s, are optimized with HpBandSter Sanchez et al (2021).

|  | Method | Summary statistics | ABC correction | Validation MSE | Test MSE |
| --- | --- | --- | --- | --- | --- |
| 3 | ABC | Yes | Single layer NN | 0.352 | 0.364 |
| 4 | MLP | Yes | No | 0.399 | 0.437 |
| 5 | MLP | No | No | 0.690 | 0.675 |
| 6 | Custom CNN | No | No | 0.485 | 0.487 |
| 8 | Flagel CNN 0/-1 encoding | No | No | 0.437 | 0.444 |
| 11 | SPIDNA | No | No | 0.453 | 0.454 |
| 29 | ABC + SPIDNA | Yes | Linear reg. | 0.335 | 0.347 |
| 32 | Attentive-SPIDNA | No | No | 0.314 | 0.320 |
| 33 | Attentive-SPIDNA with attention on scenarios | No | No | 0.288 | 0.287 |
| 34 | Attentive-SPIDNA with attention on scenarios unfreezing | No | No | 0.234 | 0.251 |

To quantify the contribution of each component of our new architecture, we performed an ablation study evaluating three versions of Attentive-SPIDNA (indexes 32, 33 and 34 in Table 1). The first version, which excludes attention over scenario replicates, reached an error of 0.320 (index 32). The second version greatly improved the prediction error (0.287, indexes 33) by including an attention mechanism on the predictions of all replicates for each scenario. During this run, the architecture is first trained to perform predictions replicate wise, then its weights are frozen, while a group of layers allowing for predictions scenario wise is added and trained. Finally, the third version improved the predictions further (0.251, index 34) by not freezing the weights and training them together with the added layers (see section Optimization strategies of Attentive-SPIDNA for additional information about the weight unfreezing procedure). These three architectures are the first that do not rely on any summary statistic or ABC step to have substantially outperformed all the other methods presented here, reducing the error by up to 27.7% compared to the previous state of the art on this task (ABC combined with SPIDNA, 0.347, index 29).

### Optimizing and evaluating SPIDNA and Attentive-SPIDNA for human populations

A narrower simulation prior, tailored to the species’ history and biology, can allow us to incorporate more knowledge about the studied species and helps mitigate errors arising from discrepancies between simulated and real data. For this reason, we simulated a second dataset with priors that are more realistic for human populations by accounting for the known mutation rate, the known recombination rate distribution and parametrized growth rate distributions (see section Simulated HGDP dataset). Sampling sizes were picked to mimic the characteristics of the HGDP dataset (between 6 and 46 individuals). This second simulated dataset (denoted as HGDP simulated dataset) allows for training our methods to perform the best prediction over the real HGDP dataset with respect to the approximations of the evolutionary simulator and our a priori knowledge about human demography that are included in the priors.

We investigate the performance of a subset of architectures on this second set. Precisely, using this dataset, we train the best SPIDNA architecture (without ABC nor summary statistics, index 11 in Table 1), and all Attentive-SPIDNA version (indexes 32, 33 and 34 in Table 1) to solve the same task (reconstructing effective population sizes through time). We present the results in Table 2.

**Table 2:** Prediction errors on the simulated HGDP dataset. The SPIDNA version included here is based the same as index 11 in Table 1. Section Inference by scenario details the differences between prediction aggregators indicated in the column “scenario prediction aggregation”. Section Optimization strategies of Attentive-SPIDNA explains the differences between the training procedures indicated in the column “Training procedure”.

|  | Method | Scenario prediction aggregation | Training procedure | Validation MSE | Test MSE |
| --- | --- | --- | --- | --- | --- |
| 0 | SPIDNA | Mean | Normal | 0.554 | 0.560 |
| 1 | A-SPIDNA | Mean | Normal | 0.480 | 0.484 |
| 2 | A-SPIDNA | Attention mechanism | Normal | 0.464 | 0.468 |
| 3 | **A-SPIDNA** | **Attention mechanism** | **Unfreezing** | **0.376** | **0.392** |

It is important to note that summary statistics cannot be easily compared directly across scenarios of this dataset. Indeed, each example has a varying number of haplotypes with the purpose of representing real data heterogeneity (where a different population might have a different number of sequenced individuals). This heterogeneity impacts most summary statistics, and one would need to redesign them in order to be insensitive to the sampling size. Previous works have instead decided to simulate datasets with many individuals, recompute summary statistics on subsets corresponding to each smaller sampling size, and use different reference tables for each prediction (Boitard et al, 2016). Although this solves the issue, it is not the most efficient fashion, particularly for machine learning methods that would need to be retrained on each reference table (i.e., for each sampling size, i.e., almost each population), hence losing the advantage of a single training and fast predictions.

For these reasons, we built architectures flexible to the input size and investigated the best procedure for training them on data heterogeneous in sample size (between 10 and 100 haplotypes for each matrix, See Section Comparison of Attentive-SPIDNA batch formats and Figure S4). Thanks to this experiment, we chose to format the batches by subsampling the number of haplotypes to 50 when greater than this value in the unprocessed SNP matrix. On the other hand, matrices with fewer than 50 haplotypes are padded to reach 50 rows.

Overall, the rank between architectures is the same as in Table 1, with the different versions of Attentive-SPIDNA, (indices 1 to 3) outperforming SPIDNA by respectively 12.6%, 16.4%, and 30%. However, the errors are greater on the simulated HGDP dataset than on the previously simulated benchmark dataset, indicating that this combination of task, priors and varying number of haplotypes yield more difficult inference task. This explains why the gap in terms of prediction error between SPIDNA on the simulated benchmark dataset (0.454, index 11 in Table 1) and HGDP dataset is important (0.560, index 0 in Table 2). Similarly, all versions of Attentive-SPIDNA have a greater error on the simulated HGDP dataset (0.484, 0.468, 0.392 in Table 2, versus 0.320, 0.287 and 0.251 in Table 1). Because it has the smallest error overall, we later applied Attentive-SPIDNA with attention mechanism on scenario and weight unfreezing on predifined scenarios and the real HGDP dataset.

### Results on predefined scenarios

The performances of A-SPIDNA without attention on scenarios and trained with the “benchmark” dataset are illustrated on a subset of cattle demographic scenarios (Figure 1) that were previously investigated by Boitard et al (2016); Sanchez et al (2021). Six scenarios were simulated: “Medium”, “Large”, “Decline”, “Expansion”, “Bottleneck” and “Zigzag”. Overall, A-SPIDNA performed better than SPIDNA (Sanchez et al, 2021) by including the true effective size in almost all box plot of each scenario. Compared to the methods developed in Sanchez et al (2021), A-SPIDNA was able to reconstruct the “Bottleneck” scenario and the most ancient bottleneck of the “Zigzag” scenario (Figure 1) with a good time estimate of both events. However, similarly to SPIDNA, predictions of very recent effective population sizes were slightly biased toward the center of the prior distribution.

**Fig. 1:**
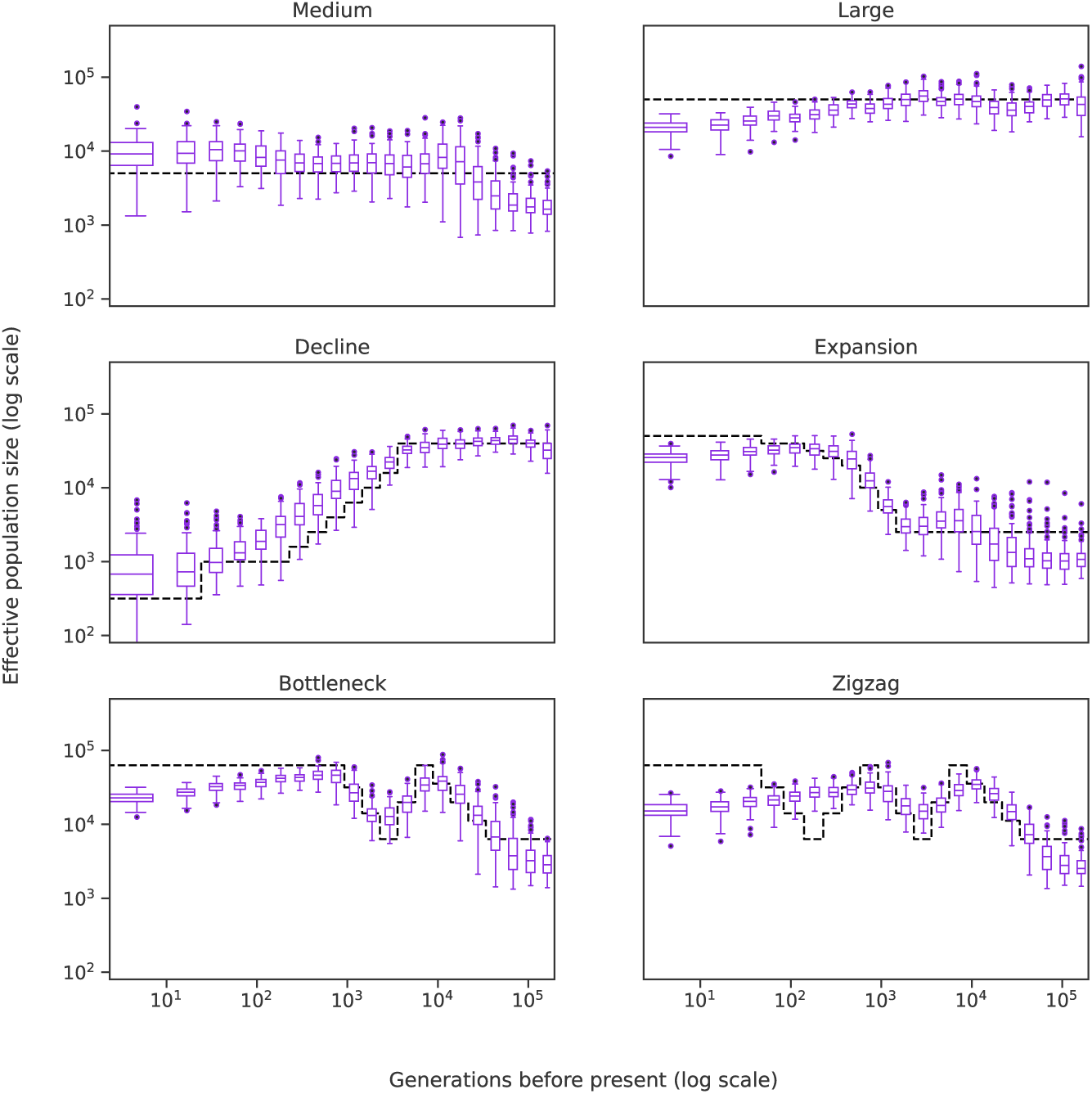
Predictions of A-SPIDNA trained on the “benchmark” dataset, for six predefined scenarios (dashed black lines). 100 replicates were simulated for each scenario. Boxplots show the dispersion of A-SPIDNA predictions over replicates.

### Reconstructing effective population size histories using the Human Genome Diversity Project

The results of our previous experiments on the simulated “benchmark” test set and simulated HGDP test set, presented in Tables 1 and 2, show that all versions of Attentive-SPIDNA have outperformed the other architectures in terms of prediction error. We selected the best architecture configuration and batch format trained on the simulated HGDP data (index 3 in Table 2) to finally infer the effective population size history of the 54 HGDP populations from the 929 whole genome sequences assuming panmixia within each population. The results presented in Figures 2 and 3 bring insights on the complex demographic processes involved in shaping observed extant human genome-wide diversity patterns at a worldwide scale.

**Fig. 2:**
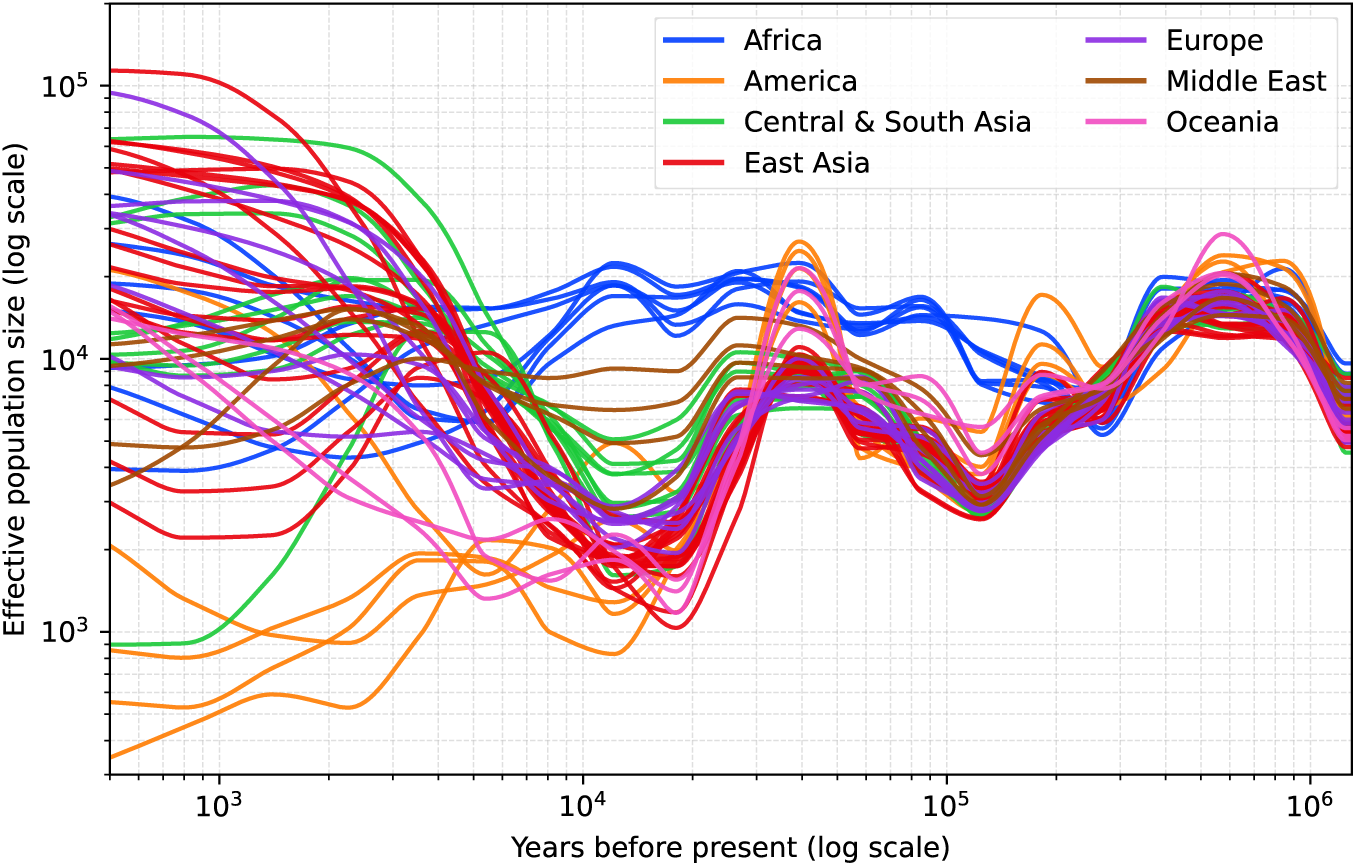
Effective population sizes inferred by Attentive-SPIDNA for the human populations from the HGDP dataset. Effective sizes are inferred by the Attentive-SPIDNA version with the lowest prediction error on the simulated dataset, i.e., with attention mechanism on scenario, batches with padding and the weight unfreezing mechanism. For each population, the 21 predicted values are interpolated using a piecewise cubic Hermite interpolator.

**Fig. 3:**
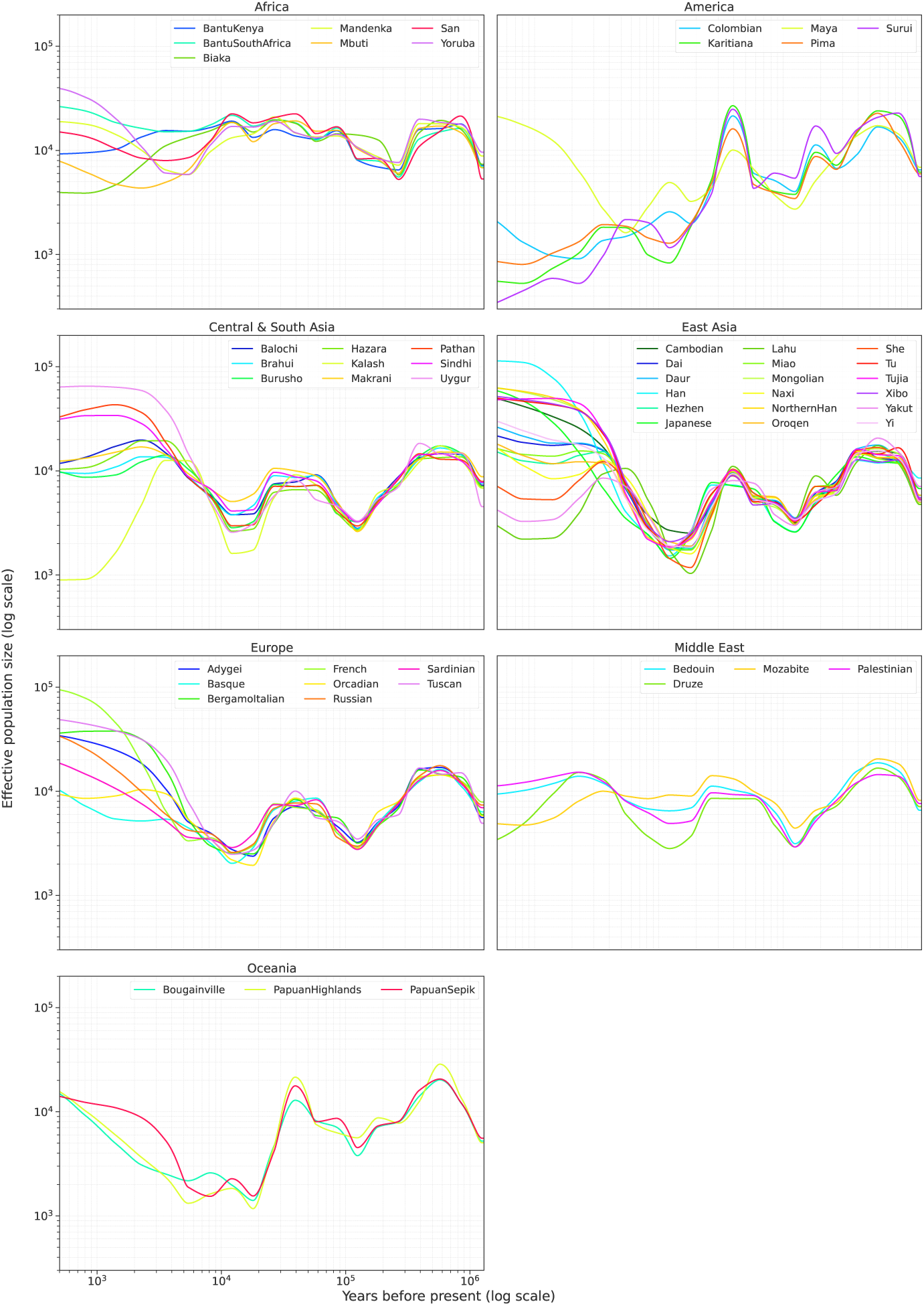
Effective population sizes inferred by Attentive-SPIDNA for the human populations from the HGDP dataset separated by region. Effective sizes are inferred by the Attentive-SPIDNA version with the lowest prediction error on the simulated dataset, i.e., with attention mechanism on scenario, batches with padding and the weight unfreezing mechanism. For each population, the 21 predicted values are interpolated using a piecewise cubic Hermite interpolator.

As expected due to the common origin of our species, Figure 2 shows that the inferred demographic histories of human populations gradually diverge when moving forward to the present time. Interestingly, we find ancient decreases of all population’s demographic histories between roughly 500,000 and 250,000 yBP, at which point, forward in time, the demographic history of all African populations started to diverge from that of all non-African populations. Indeed, all African populations increased in size at a relatively slow rate between 250,000 and 70,000 yBP. Conversely, between roughly 250,000 and 120,000 yBP non-African populations continued to decrease in effective sizes, before all experiencing similar increases until roughly 70,000 yBP when the demographic histories start to substantially diverge between populations. These ancient demographic patterns of our species are generally very consistent with previous work using a variety of classical simulation-based maximum-likelihood or ABC approaches relying on sets of population genetics’ classical summary-statistics computed on a variety of genome-wide data in similar or analogous population sample sets (Mallick et al, 2016; Bergström et al, 2020; Lorente-Galdos et al, 2019; Ragsdale et al, 2023; Lorente-Galdos et al, 2019; Fan et al, 2023). The onset of the most ancient decline likely reflects the speciation period of *Homo sapiens* until roughly 300-250,000 yBP. Then, ancient complex population structure in Africa, with or without the additional hypothesis of gene-flow from ancient non-*Homo sapiens* lineages (Lachance et al, 2012; Lorente-Galdos et al, 2019; Ragsdale et al, 2023; Fan et al, 2023; Breton et al, 2024), explains consistently our results for the onset of the diverging demographic patterns, i) among African populations, and ii) between African populations and the group of populations that would latter become, roughly 150,000 yBP, non-African populations after the Out-Of-Africa.

After the Out-of-Africa roughly 120,000 yBP, our results recapitulate general expectations from the sequence of serial-founder events of peopling of the Middle East, Central and South Asia, Europe, East Asia, Oceania, and the Americas for the HGDP population set (e.g. Ramachandran et al (2005); Bergström et al (2020)). Indeed, Figure 3 explicitly shows between 120,000 and 70,000 yBP, the first decrease in population size followed by an increase at a high rate due to the initial founding event Out-of-Africa into the Middle East and Central Asia synchronic for all non-African populations. Then we found another strong decline in population sizes between roughly 60,000 and 20,000 yBP respectively in Europe and East Asia also consistent with another series of founding events for the separate peopling of either region from the Middle East and Central Asia. Finally, note that these decreases continue for most American and Oceanian populations until more recently (roughly 15 to 5,000 yBP) indicating the last series of founding events from East Asia separately into these two last major geographic regions settled anew by *Homo sapiens*.

Figure 3 shows the increasing variation in the demographic histories of worldwide populations during the last roughly 10,000 years until present, as previously observed in numerous studies, including in Bergström et al (2020). Interestingly, while, regionally, certain outlying demographic patterns during this period, such as for the Central Asian Kalash or the American Maya, are also found in Bergstrom et al.’s SMC++ approach, the results obtained here with A-SPIDNA markedly differ from this previous study concerning hunter-gatherer populations from Central (Mbuti and Biaka), and Southern (San) Africa. Indeed, Bergstrom et al. in general found marked demographic regimes since the last 10,000 years between most agriculturalist and most hunter-gatherer populations, with the former experiencing rapid growth in the last 5,000 years while the latter experienced strong declines. Here, we found less marked differences between the two groups of populations even if our inferences also result in relatively smaller effective sizes in hunter-gatherer populations compared to agriculturalists in the present. Nevertheless, note that our posterior estimates are largely comparable with several other previous investigations also using estimation approaches relying on different model and simulation specifications, based on ML or ABC, (e.g. Lachance et al (2012); Lorente-Galdos et al (2019); Ragsdale et al (2023); Fan et al (2023); Breton et al (2024)). Since our A-SPIDNA approach captures fundamentally different aspects of genomic information compared to these previous approaches, our results illustrate that A-SPIDNA will significantly contribute to provide future novel insights into the highly complex evolutionary history of human populations.

Finally, we do not observe a correlation between the number of sampled haplotypes and the inferred effective population sizes (Figure S9). To confirm this, we downsampled the number of haplotypes within HGDP populations and reiterated the inference. This showed no substantial differences due to sampling size except when analyzing as few as 5 haplotypes, which is outside both our training prior distribution and real data range (Figure S10).

## Materials and methods

### Simulated and real datasets

To evaluate the capability of our deep learning approach in handling genomic data for population genetics, we focused on a complex task of demographic inference. Specifically, our goal was to predict effective population size values across multiple time windows within a specified single population model. To this end, we simulated two datasets of whole-genome population samples from two different scenario priors. These datasets were used to train and benchmark the performance of A-SPIDNA against other approaches. Subsequently, we assessed the performance of A-SPIDNA on a real dataset comprising full human genomes from the Human Genome Diversity Project (HGDP) (Bergström et al, 2020).

### Data format

We used the *SNP matrix* data format as described in Sanchez et al (2021). A sample of multiple individuals from the same population is represented by a SNP matrix and its associated position vector encoded as distances (in bp) between SNPs. The matrices contain zeros and ones to encode ancestral and derived alleles or minor and major frequency alleles, depending on the information available for the real dataset. All datasets used here are phased. Section 1.2 explains in details the different methods tested to concatenate SNP matrices into batches.

### Real data from the HGDP dataset

The HGDP dataset (Bergström et al, 2020) comprises 929 whole genomes from 54 populations, with 6 to 46 individuals per population. Sequencing was performed using Illumina technology with an average coverage of 35x and reads have been mapped to the GRCh38 reference assembly. Although the HGDP dataset includes 1575 fewer sequenced genomes than the 1000 Genomes Project, Bergström et al (2020) were able to identify a number of SNPs of the same order thanks to their high-coverage and the diversity of their sample. For our analysis, we used the statistically phased version of the dataset, which allowed us to conserve the haplotype information by encoding two rows per individual in the SNP matrix. This statistical phasing was compared to physical phasing by Bergström et al (2020) on a subsample, revealing a maximum switch error rate of 1.2% after excluding singletons.

We processed this dataset as follows: after removing telomeres and centromeres, autosomes have been split into 2 Mbp-long segments. Then, polyallelic sites have been removed and SNPs have been encoded with zeros for ancestral and ones for derived by comparing either to the Ensembl database; or to chimpanzee reference genome; or encoded as minor and major alleles when ancestral information was not available. The number of SNPs available for each population depends on the genomic diversity and the number of sampled individuals (see Figures S6 and S5).

### “Benchmark” dataset

To compare all 34 methods, we employed the same dataset as Sanchez et al (2021), which was simulated according to the scheme developed by Boitard et al (2016). The simulation scenarios are divided into 21 time windows, with corresponding effective population sizes drawn from a log-uniform distribution. Each scenario was used to generate 100 independent 2 Mbp-long genomic segments, each consisting of 50 haploid individuals with over 400 SNPs. The complete dataset encompasses 17,961 scenarios for training, 500 for validation, and 767 for testing which makes a total of 1,922,800 SNP matrices.

### Simulated HGDP dataset

We performed simulations designed to encompass the HGDP dataset using msprime (Baumdicker et al, 2022). The 21 time windows follow the same formula as the benchmark dataset, with the oldest window representing the time before 1 million years ago. For each of 30,000 scenarios, we generated 100 segments of 2 Mbp-long regions. The mutation rate was set to 0.5 × 10*^−^*^9^ mutation per base and year, and a generation length was set to 29 years (Scally, 2016). The recombination rate *ρ*, in centimorgan per base, was randomly drawn from a kernel Gaussian distribution fitted over the distribution from the recombination map of the 1000 genomes project (Consortium et al, 2015) (see Figure S7).

We selected this recombination rate prior over a constant value or a simpler uniform distribution to more accurately account for this confounding factor. This approach gives a good balance between realism and computational feasibility, compared to the most realistic strategy that would be to simulate the complete genome alignments with corresponding recombination rates along each segment. We preferred to simulate only 100 segments of 2 Mbp-long segments per scenario, rather than simulating as many 2 Mbp-long segments as there are in a real genome, to generate a wider range of demographic scenarios. This approach allows us to focus primarily on the inference of population size histories while accounting for unknown recombination.

Contrary to the “Benchmark” dataset, the number of haplotypes varied across scenarios and was sampled was uniformly drawn between 10 and 100. The effective population size for the most ancient of the 21 time steps was drawn uniformly on a *log*_10_ scale between 100 and 1,000,000. Then, for each time window *t*, a growth rate *g*^(*t*)^ was drawn using the following formula:

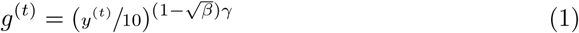

with *β* ∼ *U* (0, 1), *γ* ∼ *B*(−1, 1) and *y*^(*t*)^ the length of time window *t* in years. The growth rate is redrawn if the resulting effective population size fall outside the]100; 1, 000, 000[range. We chose this distribution to concentrate most of its mass around a growth rate of 1, while allowing for some rare extreme values depending on the duration of the time window, as shown by Figure 4. The rationale behind this choice of prior is to ensure realistic population size changes that depend on the time elapsed. This contrasts with the prior proposed by Boitard et al (2016) and Sanchez et al (2021), which limited the space of plausible histories by preventing extreme jumps (increases or decreases) in a single step, but did not account for step duration. Our approach allows for some extreme growth rates, which cannot exceed 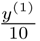.

**Fig. 4:**
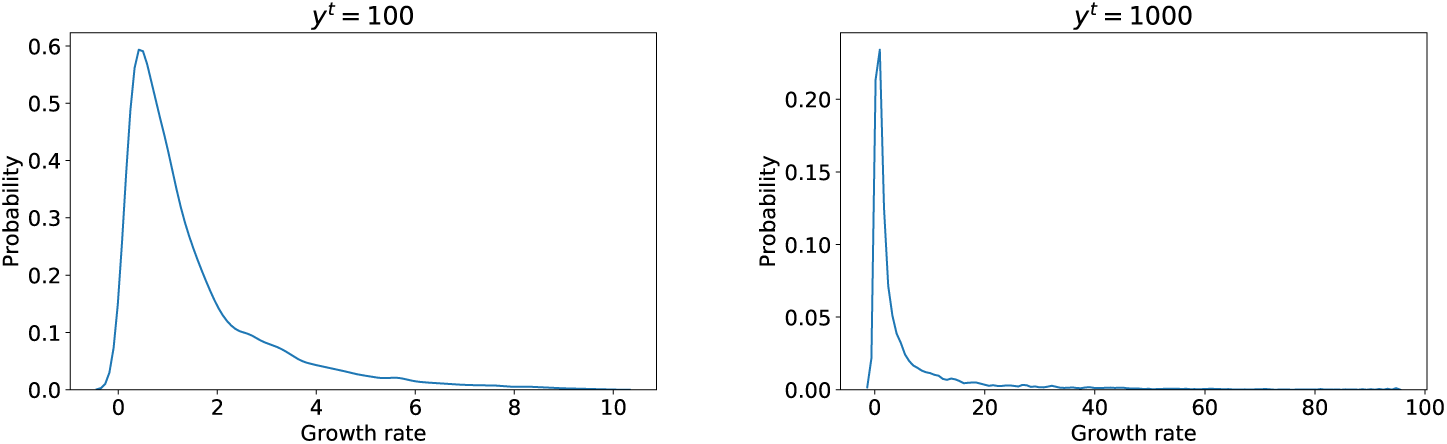
Growth rate simulation priors for time windows of 100 years (left) and 1000 years (right) in HGDP simulations. These two examples of prior distributions follow Equation 1. For each simulated demographic scenario, 21 growth rates are sampled from this distribution to generate the full scenario.

**Fig. 5:**
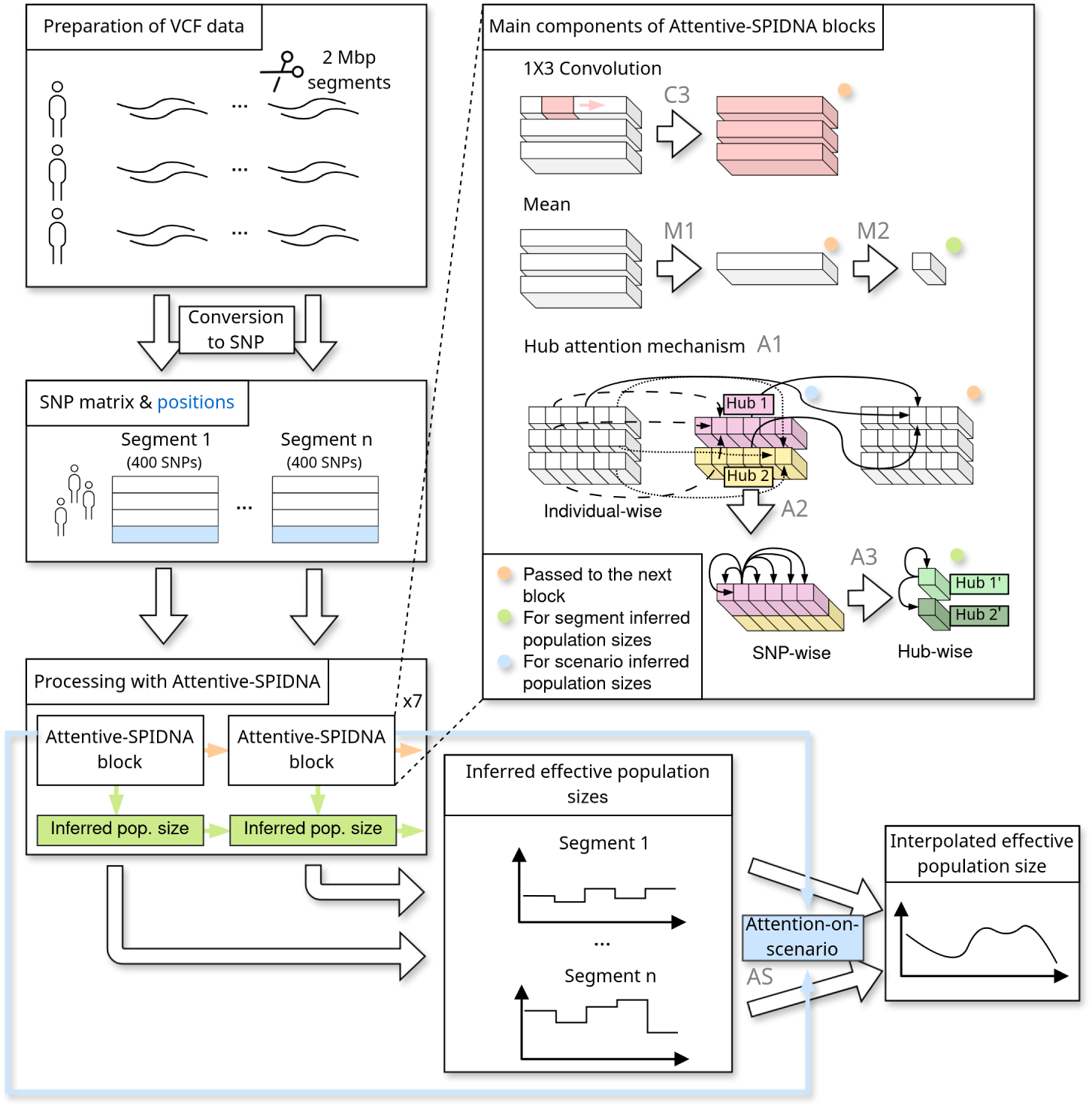
Overview of the Attentive-SPIDNA workflow and architecture. Wholegenome sequences are first divided into 2 Mbp segments, from which the first 400 SNPs of each segment are extracted to form SNP matrices to serve as input to Attentive-SPIDNA. The architecture comprises seven sequential blocks, each integrating a convolutional layer, an average layer and a hub attention mechanism (see Figure S3 for a more detailed diagram of the architecture). Finally, the inferred population sizes of each segment are aggregated using the attention-on-scenario mechanism. The labels C3, M1, M2, A1, A2, A3 and AS correspond to those shown in Figure S3.

After removing scenarios that contain at least one segment with fewer than 400 SNPs, the 21,044 scenarios are separated between a training set with 20,044 scenarios (i.e., 2,004,400 training SNP matrices) and a validation set with 1,000 scenarios (i.e., 100,000 validation SNP matrices). Independently, we simulated a test set including 1,499 scenarios (i.e., 149,900 validation SNP matrices) after preprocessing.

### Attentive-SPIDNA: a new permutation invariant ANN for demographic inference

We developed a new deep learning architecture built upon our previous work from (Sanchez et al, 2021) (as explained in section 1.1), with the primary intent of enhancing performance by addressing two of its bottlenecks using attention mechanisms. First, we improved the network’s expressivity regarding features computed across sequences by supporting the averaging layer of SPIDNA with an attention mechanism. Second, we employed another attention mechanism to better combine the network prediction made on each segment. We compared multiple versions of this architecture in order to evaluate the gains from these improvements and tested several optimization strategies.

### Improving network expressivity with the attention hub mechanism

The attention mechanism we introduced is a variation of the original self-attention mechanism from Vaswani et al (2017); that relies on a new component called hub. In the original self-attention, an affinity is computed between every pair of input elements, which leads to a *n*^2^ term in the overall complexity (where *n* is the number of haplotypes). Here instead, the affinity is computed between each input element and a predefined number of hubs, to reduce the complexity (now linear in *n*). Intuitively, these affinity values will tell how to map the values computed from each input element to the hubs. The hubs are then mixed together and mapped back to the input space thanks to another set of affinity values. In the context of a network that has SNP matrices as input, each hub is a combination of the haplotypes and expresses a specific, learnable statistic over them. The attention mechanism allows expressing statistics over individuals that share a particular genetic pattern only. section 1.1 explains in more details the attention hub mechanism.

### Combining segment predictions using attention

We later improved A-SPIDNA by adding a similar attention mechanism to combine the outputs from the different segments of one scenario. The predictions from the different segments have been previously combined simply by performing a mean, which does not take into account that different segments can contain more or less information about the demography. This attention mechanism allows the network to express how confident it is in the prediction yielded by each segment, in order to better combine them. This last iteration of A-SPIDNA being more difficult to train, we first initialized the rest of the network with a pretrained regular A-SPIDNA and optimized only the last layers (the ones that combine segments) as explained in the next section (see Figure 6). We then unfreeze all weights to fine-tune the whole network. Details about the implementation of this attention mechanism can be found in Supplementary Text, section 1.1

**Fig. 6:**
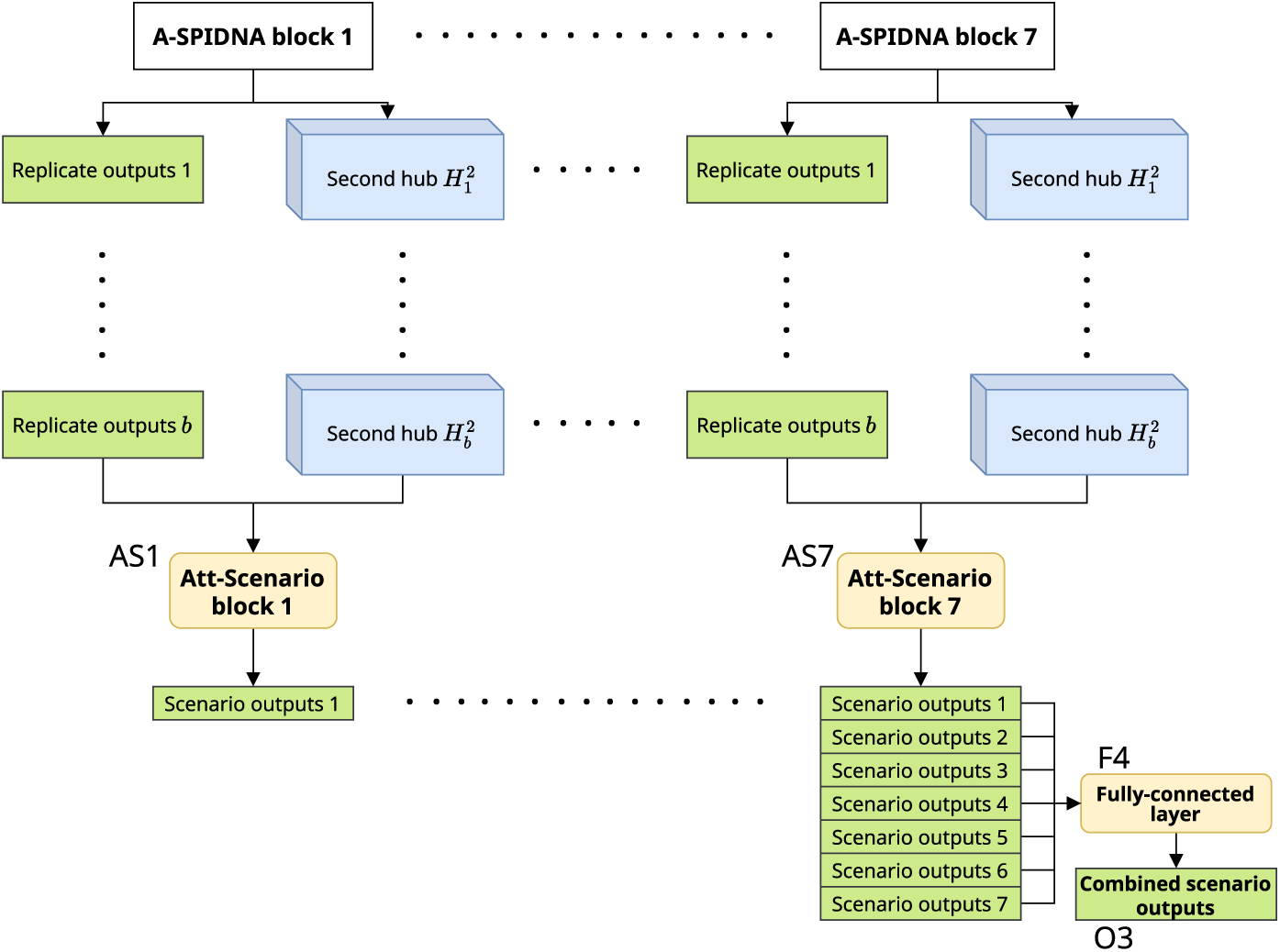
Each block of the Attentive-SPIDNA architecture outputs 21 demographic parameter values and a tensor of hub values. The attention on scenario then combines the demographic parameter values from all the replicates using weights determined by the hub values. The 7 vectors of 21 demographic parameter values are finally combined into one using a fully-connected layer. The labels AS1, AS7, F4 and O3 follow the same labeling principle as in Figure S3.

### Optimization strategies of Attentive-SPIDNA

We tested numerous versions of A-SPIDNA during its development, but we only focus on three here: A-SPIDNA, A-SPIDNA with attention on scenario and A-SPIDNA with attention on scenario unfreezing. All three versions include a learning rate decay strategy, halving the learning rate after five epochs to improve the final prediction error. When trained on the dataset from Sanchez et al (2021), these versions of ASPIDNA use the same batch format as SPIDNA with batch normalization (batches of SNP matrices with 50 haplotypes and 400 SNPs). However, because the HGDP dataset includes varying number of haplotypes per scenario to mimic the different sample sizes of the real HGDP populations, we tested different mini-batch formatting strategies on the benchmark dataset and applied the best strategy to the HGDP dataset. Both A-SPIDNA with attention on scenario and A-SPIDNA with attention on scenario unfreezing are trained in two phases. First they are trained to predict demographic parameters segment-wise, like A-SPIDNA, for ten epochs. Then, they are trained to predict demographic parameters scenario-wise, using all segments of a same scenario. A-SPIDNA with attention on scenario optimizes only the weights and biases used to combine segment predictions into one scenario prediction during this second phase. In contrast, A-SPIDNA with attention on scenario unfreezing optimizes all parameters, but assigns a higher learning rate to the weights and biases used for combining the predictions on all segments. This approach is intended to stabilize learning when transitioning from segment-wise to scenario-wise prediction task. In experiments not shown here, training scenario-wise A-SPIDNA without pretraining on segments resulted in unstable behaviours, with increasing error on the training set.

## Discussion

Identifying effective ways to process multiple DNA sequences in a deep learning framework remains an open challenge, that is not only limited to demographic inference, but to a broad range of genomics applications. In this work, we followed a set of guiding principles to inform our architectural choices, leading to a substantial improvement over state-of-the-art methods for demographic inference. Rather than relying solely on the network’s capacity to implicitly learn the structure of genomic data, we explicitly incorporated data characteristics into the model design, drawing inspiration from both summary statistics commonly used in population genetics and recent advances in deep learning architectures. Attentive-SPIDNA combines convolutional layers and an attention hub mechanism to capture inner structures, local and longrange dependencies analogous to underlying genealogical relationships and linkage disequilibrium, while avoiding the quadratic complexity of standard self-attention. The architecture leverages multiple levels of variability in the data, including SNP values, inter-SNP distances, haplotype-level heterogeneity, variable sample sizes, and region-to-region variation in signal strength, while remaining invariant to haplotype ordering and flexible to heterogeneous genomic datasets. By introducing attention hubs as a richer alternative to simple invariant means over haplotypes, together with an attention-based mechanism for aggregating information across genomic regions, Attentive-SPIDNA is the first fully end-to-end approach to outperform ABC-based methods on this benchmark. Moreover, its successful application to HGDP data further demonstrates its practical relevance on real genomes.

The performance improvement of Attentive-SPIDNA compared to SPIDNA can be attributed mostly to the addition of the attention hub mechanism and the attention over scenarios, as shown by the ablation study. In SPIDNA, and other exchangeable networks, permutation invariance is enforced through a simple averaging aggregation operators over haplotypes, which can act as an information bottleneck by collapsing individual-level variation into a single global descriptor. Although such mean-based architectures are theoretically universal in a Deep sets architecture, the complexity and depth required to approximate informative invariant functions in practice remain unknown. By contrast, attention hubs allow learning statistics over subsets of haplotypes, enabling differential contributions of individuals depending on their genetic patterns. Adding these mechanisms aligns naturally with population genetics intuition, where demographic information is often unevenly distributed across haplotypes and genomic regions, or subsets of closely related individuals carrying disproportionate signal. The network constructs optimized multi-individual and genome-wide features that provide alternative information compared to, e.g., pairwise sequence comparisons done in PSMC-like approaches. Within a genomic region, our approach shares similarities with MSA Transformers (Rao et al, 2021), which were originally designed for non-population-genetics applications. However, we focus exclusively on row-wise attention (across sequences) and complement it with convolutional operations along the SNP dimension to efficiently capture local patterns. Unlike the MSA Transformer, our attention mechanism scales linearly with the number of haplotypes, avoiding quadratic complexity. A similar strategy, using global tokens to reduce sequence-length complexity, was previously introduced for single sequences in BigBird (Zaheer et al, 2020). Recently, Korfmann et al (2026) have leveraged attention mechanisms (through a language model) to accurately estimate pairwise coalescent times and their uncertainty along the genome. In their case, attention is applied across genetic markers. Overall, attention-based invariant representations combined with local 1D convolutions offer a more efficient and expressive alternative to simple averaging, while preserving permutation invariance and scalability to variable sample sizes. By efficiently reusing learnable parameters, these mechanisms also allow the network to remain relatively compact and fit on a single GPU, making it easier for the community to retrain and use.

Despite these advances, our approach remains constrained by the scarcity of real, annotated genomic datasets and therefore relies on how closely the evolutionary models used for simulation approximate the true evolutionary process. As with other simulation-based inference methods, this dependency on the simulator represents an important source of uncertainty. While likelihood-based methods also depend on correct evolutionary models and are most often not applicable to complex scenarios, our approach offers a key advantage: the inference stage is decoupled from data simulation, making it straightforward to incorporate new hypotheses about evolutionary processes by modifying the simulations, while retaining a similar deep learning architecture for inference. Based on test sets, the user can evaluate the inference capacity of the network itself, as well as the performance loss under a misspecified model. When applied to real data, unexpected predictions should raise concern and foster changes in priors and modeling assumptions. This can be achieved in a more principled manner. Our study focused on improving the expressiveness of frugal exchangeable networks, which can now be integrated with (i) estimating posterior distributions to quantify parameter uncertainty (Chan et al, 2018; Sanchez et al, 2021; Min et al, 2025; Korfmann et al, 2026), (ii) approaches estimating both this uncertainty and the epistemic uncertainty informative of the gap between training set modeling assumptions and reality (Cury et al, 2022), and (iii) domain adaptation strategies reducing this gap (Mo and Siepel, 2023; Cobb and Smith, 2025). More generally, the use of deep learning offers great flexibility with respect to inference objectives, as the same framework can be retrained to target different evolutionary parameters, such as introgression, selection, population structure and dispersal, by adapting the simulated data and network outputs. In this study, we investigated the inference of 21 effective population size values corresponding to the simulator time windows, in order to facilitate comparison with previous works (Boitard et al, 2016; Sanchez et al, 2021). Nonetheless, the flexibility of the approach allows extension to other evolutionary models and alternative sets of parameters, as demonstrated by Korfmann et al (2024).

In conclusion, Attentive-SPIDNA demonstrates that attention mechanisms are valuable for processing samples of DNA sequences and addressing challenging inference tasks such as reconstructing demographic histories from genomic data. By explicitly integrating key characteristics of DNA samples into the network design through attention-based components, this framework provides a general and effective strategy to extract information embedded into multiple DNA sequence alignment that can be extended to other inference problems relying on similar data. Moreover, the flexibility of deep learning architectures makes such approaches well suited to adapt to future advances in the field without requiring fundamental modifications to the overall framework.

## Data and code availability

Attentive-SPIDNA is available in the dnadna package (Sanchez et al, 2023). A table of A-SPIDNA inferred effective population sizes for HGDP, together with the code used for plotting and simulations, is available in the Attentive-SPIDNA repository. The benchmark is the same as in Sanchez et al (2021); the corresponding simulation code is available in the SPIDNA repository. The HGDP dataset is available from the Sanger Institute website (Bergström et al, 2020).

## Acknowledgements

We thank Samuel Pavard for discussing human demographics’ priors, the Bioinfo team and Evogenomics.ai network for general discussions, Inria TAU for GPU computing resources, ANR-20-CE45-0010-01 RoDAPoG and Human Frontier Science Project (number RGY0075/2019) for funding. EM Bray, Jean Cury, Elliot Mâıtre, Julien Rabault for contributing to the dnadna software at different stages of its development. PNRIA for supporting software development.

## 1 Supplementary materials

**Table S1:** Prediction errors of each method on the simulated benchmark dataset. All method’s hyperparameters but Attentive-SPIDNA’s are optimized with HpBandSter (Falkner et al, 2018). See Sanchez et al (2021) for an explanation of the adaptive characteristic of SPIDNA to the number of SNP indicated in column “Adaptive” and the utility of the parameter *α* in SPIDNA indicated in column “*α* value”.

| | Method | Adaptive | Summary statistics | ABC correction | $\alpha$ value | Validation MSE | Test MSE |
| --- | --- | --- | --- | --- | --- | --- | --- |
| 0 | ABC | No | Yes | No | No | 0.490 | 0.496 |
| 1 | ABC | No | Yes | Linear reg. | No | 0.357 | 0.369 |
| 2 | ABC | No | Yes | Ridge reg. | No | 0.363 | 0.376 |
| 3 | ABC | No | Yes | Single layer NN | No | 0.352 | 0.364 |
| 4 | MLP | No | Yes | No | No | 0.399 | 0.437 |
| 5 | MLP | No | No | No | No | 0.690 | 0.675 |
| 6 | Custom CNN | No | No | No | No | 0.485 | 0.487 |
| 7 | Flagel CNN 0/-1 encoding | No | No | No | No | 0.537 | 0.541 |
| 8 | Flagel CNN 0/-1 encoding | Downsampling | No | No | No | 0.437 | 0.444 |
| 9 | Flagel CNN -1/1 encoding | No | No | No | No | 0.610 | 0.609 |
| 10 | Flagel CNN -1/1 encoding | Downsampling | No | No | No | 0.482 | 0.484 |
| 11 | SPIDNA | No | No | No | No | 0.453 | 0.454 |
| 12 | SPIDNA | No | No | No | No | 0.637 | 0.641 |
| 13 | SPIDNA | Yes | No | No | No | 0.487 | 0.489 |
| 14 | SPIDNA | No | No | No | 0.849 | 0.592 | 0.599 |
| 15 | SPIDNA | Yes | No | No | 0.539 | 0.466 | 0.469 |
| 16 | ABC + SPIDNA | No | No | No | No | 0.462 | 0.462 |
| 17 | ABC + SPIDNA | No | No | Linear reg. | No | 0.364 | 0.377 |
| 18 | ABC + SPIDNA | No | No | Ridge reg. | No | 0.371 | 0.380 |
| 19 | ABC + SPIDNA | No | No | Single layer NN | No | 0.363 | 0.372 |
| 20 | ABC + SPIDNA | Yes | No | No | 0.539 | 0.458 | 0.460 |
| 21 | ABC + SPIDNA | Yes | No | Linear reg. | 0.539 | 0.363 | 0.369 |
| 22 | ABC + SPIDNA | Yes | No | Ridge reg. | 0.539 | 0.382 | 0.391 |
| 23 | ABC + SPIDNA | Yes | No | Single layer NN | 0.539 | 0.374 | 0.384 |
| 24 | ABC + SPIDNA | No | Yes | No | No | 0.476 | 0.478 |
| 25 | ABC + SPIDNA | No | Yes | Linear reg. | No | 0.339 | 0.353 |
| 26 | ABC + SPIDNA | No | Yes | Ridge reg. | No | 0.341 | 0.357 |
| 27 | ABC + SPIDNA | No | Yes | Single layer NN | No | 0.345 | 0.361 |
| 28 | ABC + SPIDNA | Yes | Yes | No | 0.539 | 0.474 | 0.478 |
| 29 | ABC + SPIDNA | Yes | Yes | Linear reg. | 0.539 | 0.335 | 0.347 |
| 30 | ABC + SPIDNA | Yes | Yes | Ridge reg. | 0.539 | 0.339 | 0.354 |
| 31 | ABC + SPIDNA | Yes | Yes | Single layer NN | 0.539 | 0.347 | 0.365 |
| 32 | Attentive-SPIDNA | No | No | No | No | 0.314 | 0.320 |
| 33 | Attentive-SPIDNA with attention on scenarios | No | No | No | No | 0.288 | 0.287 |
| 34 | Attentive-SPIDNA with attention on scenarios unfreezing | No | No | No | No | 0.234 | 0.251 |

### 1.1 Attentive-SPIDNA architecture

The A-SPIDNA architecture has been build upon the SPIDNA architecture from Sanchez et al (2021). It takes the same data format as input, has the same first layers and also uses a series of blocks that updates the outputs (demographic parameters to be estimated) before passing the data to the next block. The main differences happen inside the block (now called A-SPIDNA block) depicted in Figure S1. Thanks to the addition of an attention hub mechanism (explained in the next section), the features that are passed to the next block can now be more complex. These features of the mean M1 and attention hub A1 are mixed thanks to a fully-connected layer F3. Then, they are concatenated (I3) with the features from the convolution C3 and a max pooling layer is applied before being passed to the next block.

In parallel, the first attention hub mechanism A1 also contributes to the overall inferred values by the network. Before being projected back to the output dimensions, the first hubs are duplicated and sent to a series of two other hub attention mechanisms (A2 and A3 in Figure S1) with attention computed over the hubs. Finally, predictions from A3 are combined to the predictions from the mean over the haplotype (M1) and SNP (M2) dimensions thanks to a fully-connected layer (F2). The part of the network that outputs predictions based on the mean over the haplotype and SNP dimensions (F2) is the same has the original SPIDNA architecture, but its fully-connected layer maps all the features to the 21 outputs (previously only the first 21 features were mapped to the 21 outputs) as it has shown to be a simple improvement of the SPIDNA architecture during the development of A-SPIDNA.

**Fig. S1:**
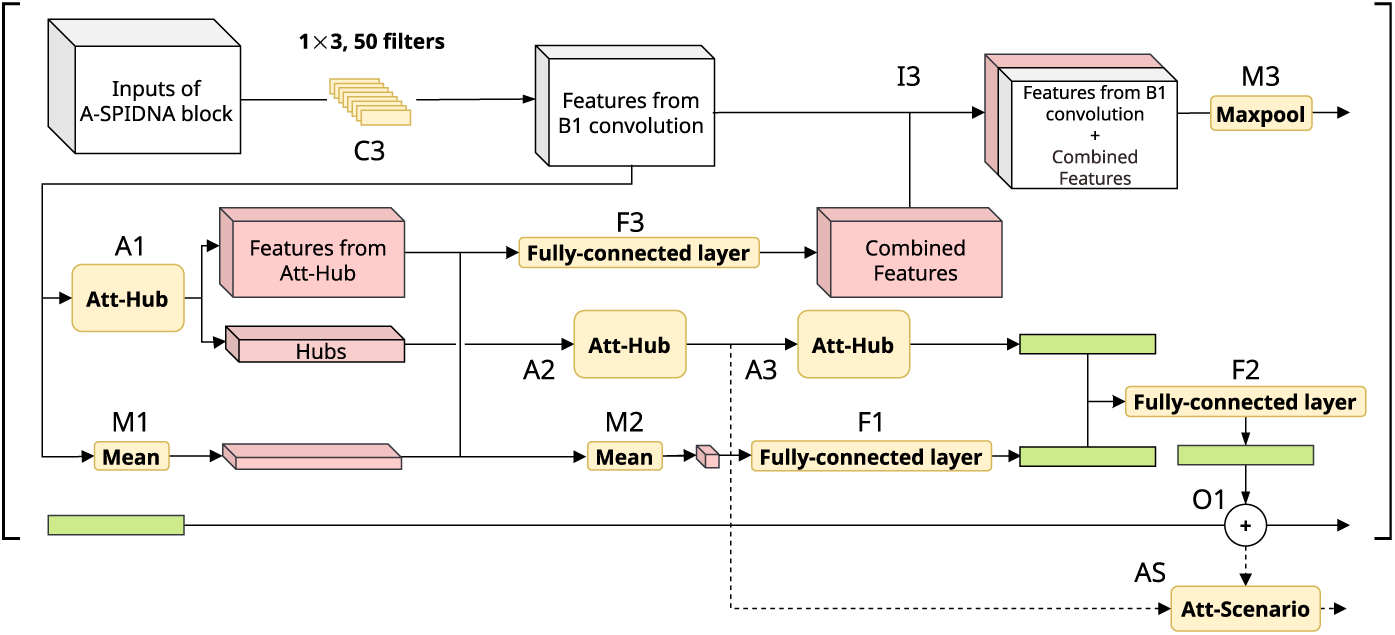
Schematic of Attentive-SPIDNA architecture block. Each A-SPIDNA block starts with a convolution layer (C3) followed by the mean over rows of the convolution layer result (M1) and the mean over columns of M1 result (M2). The output of M2 is processed by a fully-connected layer (F1) to constitute the first part of the block output. In parallel, an attention hub (A1) process C3 output. The hub features from A1 are then process by two attention hubs (A2 and A3) to constitute the second part of the block output. The fullyconnected layer F2 combine the two block output parts, and the result is added (O1) to the network output (in green) to be passed to the next block. The next block also takes as input the combination of C3 output and outputs of M1 and A1 with a fully-connected layer F3. Outputs of C3 and F3 are combined (I3) and passed to the next block after passing through a max pooling layer M3.

#### Attention hub mechanism

To express richer statistics, the attention hub mechanism is added to each SPIDNA block alongside the computation of the mean over the haplotypes’ dimension. First, a set of keys *K* and values *V* are computed with two fully-connected layers over the features dimension of the input tensor. Then a third set of affinity values *A^in^* between each element of the sequence (here the elements are a set of features corresponding to a haplotype and a SNP) and each hub is computed using a fully-connected layer with *K* as input. The outputs *A^in^* of this fully-connected layer replace the product between *Q* and *K* from the original attention mechanism (Vaswani et al, 2017). These affinity values are passed through a softmax function and multiplied to the set of value *V* to create the hubs *H*^1^. This way, each hub selects individuals, according to their descriptors *K*, and mix their values *V*, in a weighted sum depending on *K*. In our architectures, we set up the fully-connected layer dimensions so that operation yields 10 hubs, with 50 features each and as many “pseudo-SNPs” (elements of the SNP dimension) as the inputs of the attention mechanism. Each hub processes its input data internally (through 2 fully-connected layers), independently of other hubs. Hubs are passed to another fully-connected layer (*H*^2^) in order to match their dimensions with the output dimensions (*O*), in the case where the number of features computed for *K* and *V* is different from the number of features required for the outputs. The hubs then dispatch the information they computed to each individual. For this, another attention mechanism allows each individual to choose which hubs it would like to listen to (by expressing weights for each hub). Therefore, another set of affinity values *A^out^* between hub and the original input data is computed with a fully-connected layer over the inputs *X* and then passed through a softmax. In other terms, the affinities *A^out^* determine the contribution of each hub to the output. Finally, the hubs are mapped to the output space by multiplying them with this second set of affinities *A^out^*and sent to the next A-SPIDNA block. Figure S2 shows an overview of the attention hub mechanism.

**Fig. S2:**
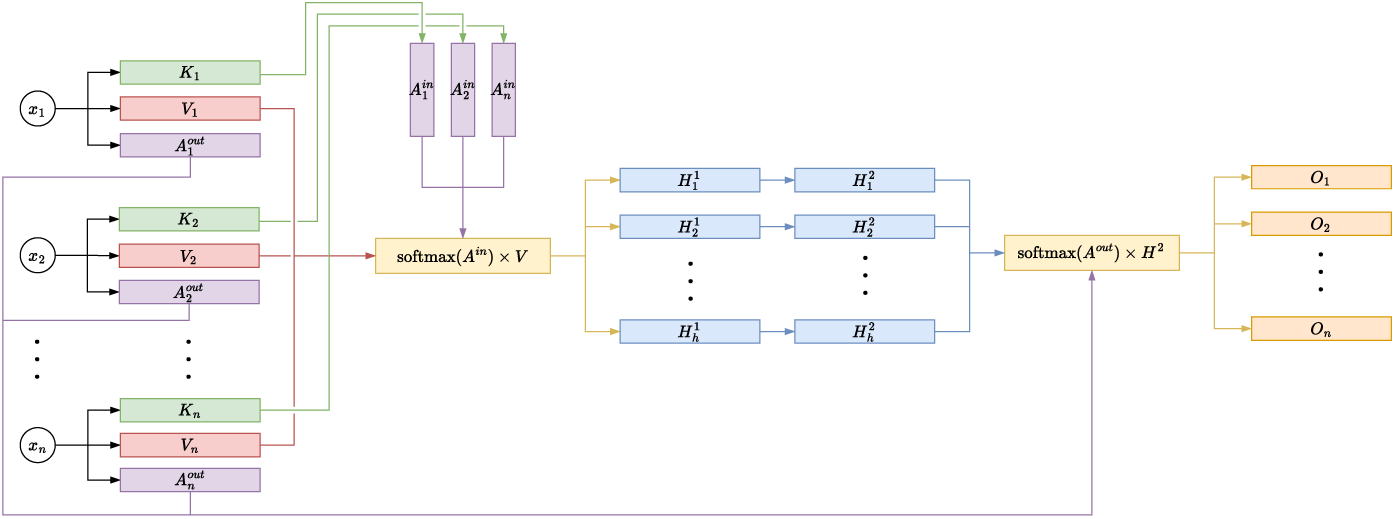
Schematic of attention hub. Two fully-connected layers compute keys *K* and values *V* from the attention hub inputs. A third fully-connected layer computes the affinity values *A^in^* from the keys *K*. The product between the softmax of *A^in^* and *V* gives the hub values *H*^1^ (softmax(*A^in^*) *× V*). A fourth fully-connected layer computes *H*^2^ from *H*^1^. The last fully-connected layer computes the affinity values *A^out^* used by the second attention dot product (softmax(*A^out^*) *× H*^2^) to output *O*.

We now consider the same dimension notation than Sanchez et al (2021) with *B* the batch dimension, *M* the row dimension (also the haplotype/genotype dimension before the first layer), *S* the column dimension (also the SNP dimension before the first layer) and *F* the feature dimension. The attention hub takes as input a tensor data of *B* × *M* × *S* × *F* and starts by swapping *M* and *S*. In practice, we choose our fully-connected layers so that values *V* have dimensions of *B* × *S* × *M* × *F*_1_ = 50, keys *K* have dimensions of *B* × *S* × *M* × *F*_2_ = 50 and affinity values *A^in^* have dimensions of *B* × *S* × *M* × *N_hubs_* = 10. After permutation of *M* and *N_hubs_* in *A^in^*, the matrix multiplication between *A^in^* and *V* gives a hub tensor *H*^1^ of *B* × *S* × *N_hubs_* = 10 × *F*_2_ = 50, transformed by a fully-connected layer into *H*^2^ of *B* × *S* × *N_hubs_* = 10 × *F_out_* = 50 dimensions. *A^out^* are computed in parallel by a fully-connected layer with the tensor data of the attention hub as input and has dimensions of *B* × *S* × *M* × *N_hubs_* = 10. The matrix multiplication between *A^out^* and *H*^2^ leads to the output tensor *O* of dimension *B* × *S* × *M* × *F_out_* = 50. Finally, *M* and *S* are swapped back so that the output dimensions correspond to the input ones.

**Fig. S3:**
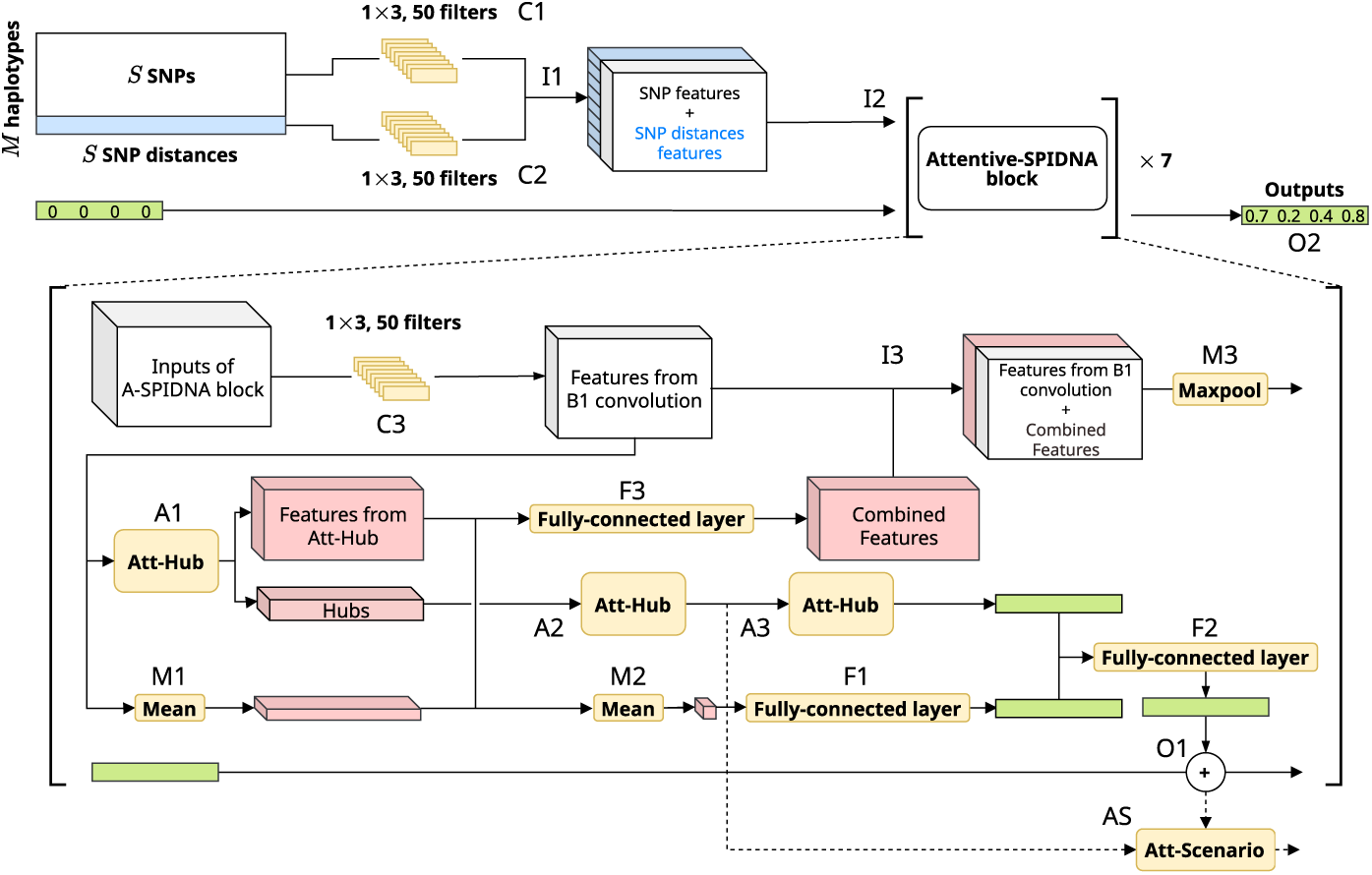
Comparison of Attentive-SPIDNA prediction error for different number of haplotypes and SNP matrix format.

#### Inference by scenario

In order to combine the predictions performed over each segment (representing one of the simulated or observed genomic regions) of one scenario (one specific population size history) in a more complex way than the mean previously used, another attention mechanism has been added to A-SPIDNA. In this version called A-SPIDNA with attention on scenario, each batch contains all segments of a single scenario and nothing more. The features computed in the second layer of hubs of each block (A2 in Figure S1) are averaged on the hub dimension and fed to a fully-connected layer followed by a softmax. The attention vector obtained determines the contribution of each segment to the final prediction by weighting the outputs (computed by F2 in Figure S1). This step finally leaves one prediction by block in the network that are combined thanks to a last fully connected layer. The idea behind this strategy is to let the network give different weights to the different segments for predicting the demographic parameters of a scenario. Therefore, the network could potentially learn to give more importance to the most informative segments.

### 1.2 Comparison of Attentive-SPIDNA batch formats

SPIDNA and Attentive-SPIDNA are designed to take a variable number of haplotypes without modification of the SNPs matrices, thanks to the use of layers that easily adapt to variations in the input sizes. More specifically, the layers that we use (the convolutional and hub attention layers) treat each element of the haplotype dimension in the same manner. Therefore, adding an element (a haplotype) to the input only requires repeating the same processing that is already applied to each other element (e.g., the filter convolutions are repeated to match the number of element of the haplotype dimension in the case of convolutional layers). Furthermore, the mean steps of the networks operate the same regardless of the number of haplotype, leading to an output of the same size when the number of haplotype varies.

However, in practice, SNP matrices with different sizes cannot be collated in the same batch (the same tensor). Therefore, training a network on these matrices would require to input them one by one and performing the backpropagation once they each have been processed by the network. This considerably reduces the parallelization of ANNs, and thus, slows down the training of the network. This is why it is preferable to collate SNP matrices in the same batch by either padding or by collating matrices of the same sizes together.

This last experiment compares four different batch formats that can be used to collate SNP matrices. To this end, between 5 and 50 haplotypes are drawn for each replicate of the simulated benchmark dataset, and SNP sites that are not polymorphic (variable) in the subset of selected haplotypes are removed. We then trained Attentive-SPIDNA by using the four batch formats described in Figure S4. After training, we evaluated the four trained architectures on the validation dataset with all matrices subsampled to between 5 and 50 haplotypes.

From Figure S4, the best method consists in making batches with SNP matrices that have a different number of haplotypes (heterogeneous batches) by padding every SNP matrices with 255 to match 50 haplotypes (blue line). We choose the value 255 because the SNP in the matrices are encoded as unsigned integer, and it is the most distinguishable value from 0 and 1 for this type of data. With this method, all the SNP matrices reach the dimension 50× 400, i.e., 50 “haplotypes” (some will be completely filled with value 255) and 400 SNPs. It shows a very constant error for any number of haplotypes with a slight increase for the smallest number, which is expected due to lack of information when the number of haplotypes is very low. Thanks to this experiment conducted on the simulated benchmark dataset, we chose the batch format later used for the simulated HGDP dataset, where matrices have a different number of haplotypes to match the different number of individuals in the real HDGP dataset.

**Fig. S4:**
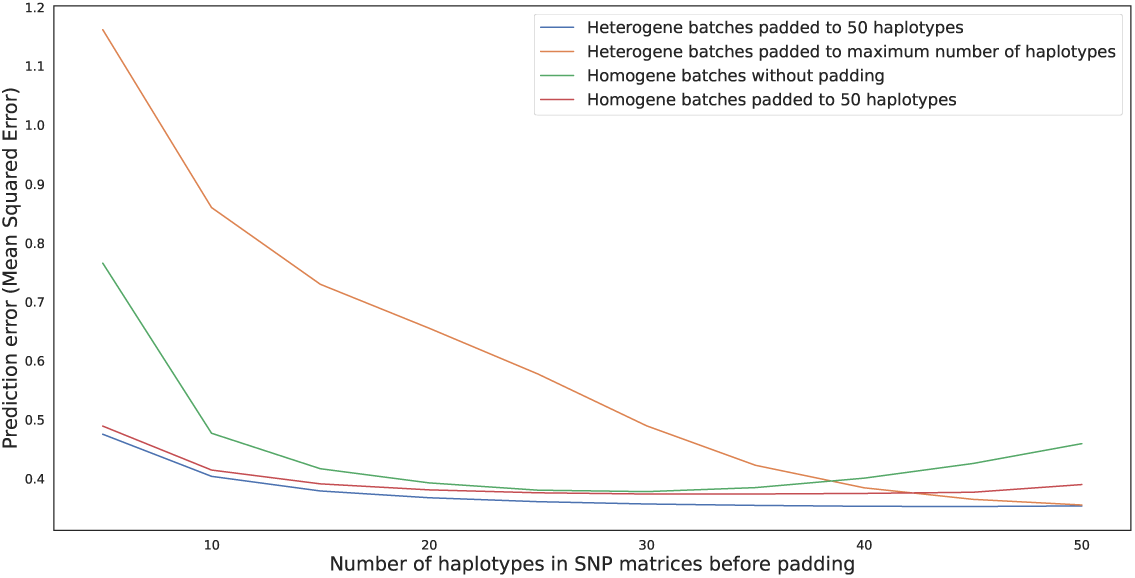
Comparison of Attentive-SPIDNA prediction error for different number of haplotypes and SNP matrix format. Each Attentive-SPIDNA architecture is trained on the simulated benchmark training dataset, with between 5 and 50 haplotypes randomly subsampled for each SNP matrix. Blue and orange lines represents Attentive-SPIDNA trained with batches composed of SNP matrices with different number of haplotypes. SNP matrices in batches are either padded with 255 to reach 50 “haplotypes” (blue line) or padded to fit the SNP matrix with the highest number of haplotypes in the batch (orange line). Green and red lines represents Attentive-SPIDNA trained with batches composed of SNP matrices with the same number of haplotypes. SNP matrices in batches are either padded with 255 to reach 50 “haplotypes” (red line) or not padded (green line). Prediction errors are computed over the benchmark validation dataset, with all matrices subsampled to the number of haplotypes indicated on the x-axis.

**Fig. S5:**
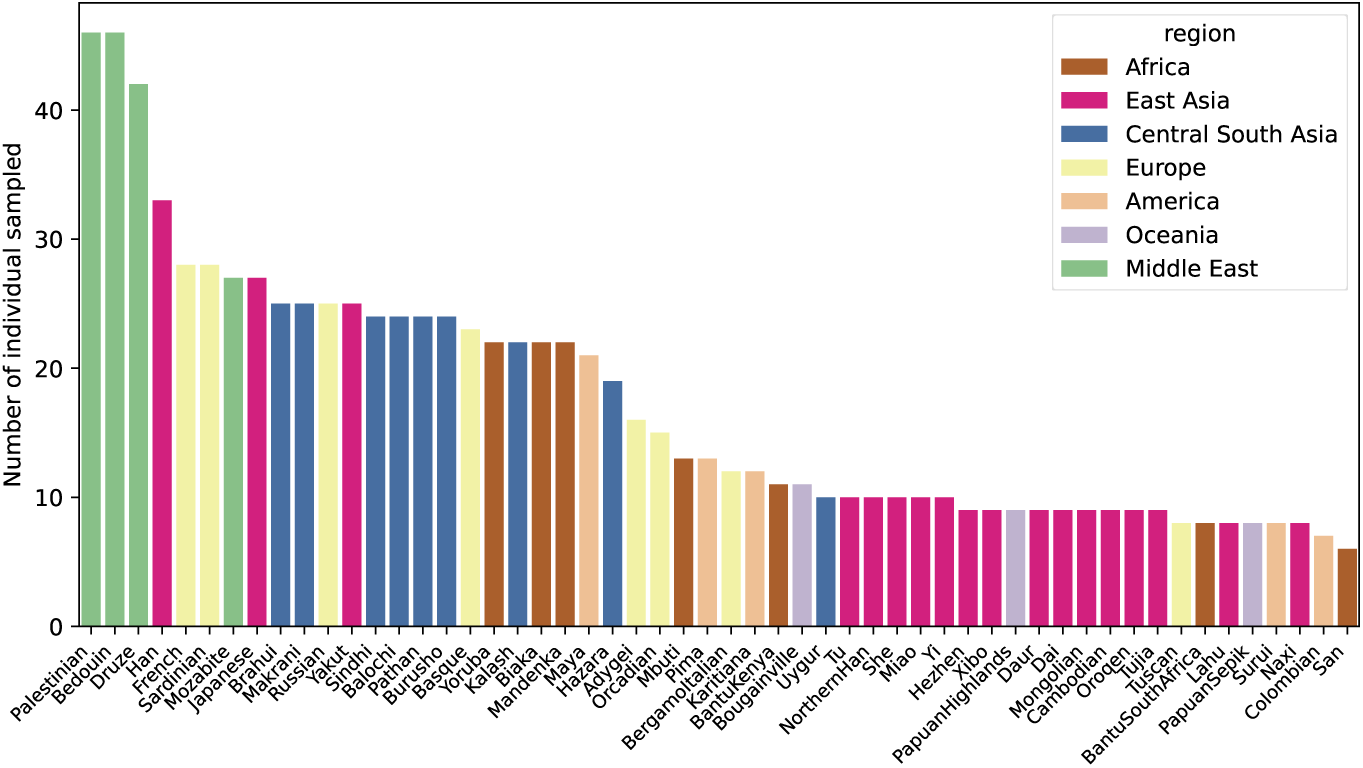
Number of sample peer population in the HGDP dataset (Bergström et al, 2020).

**Fig. S6:**
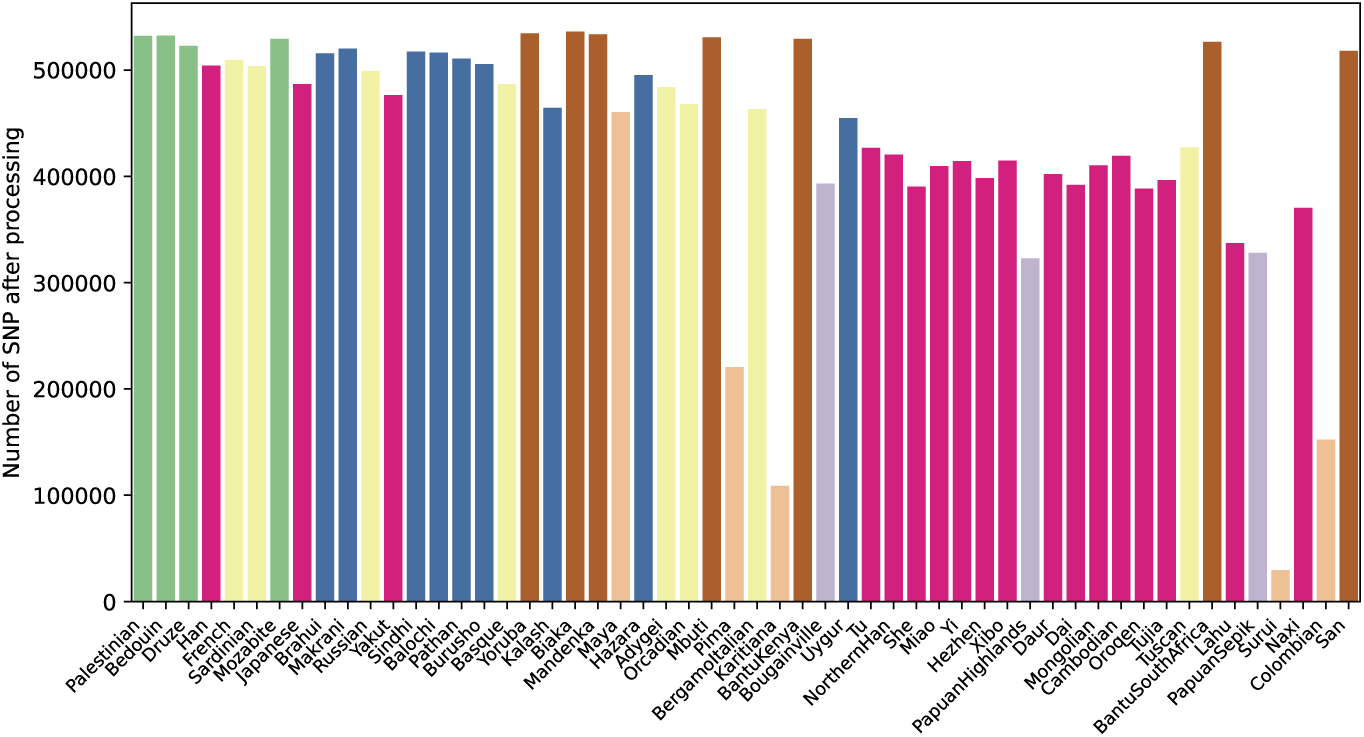
Number of SNP per population after removal of telomeres and centromeres the HGDP dataset (Bergström et al, 2020).

**Fig. S7:**
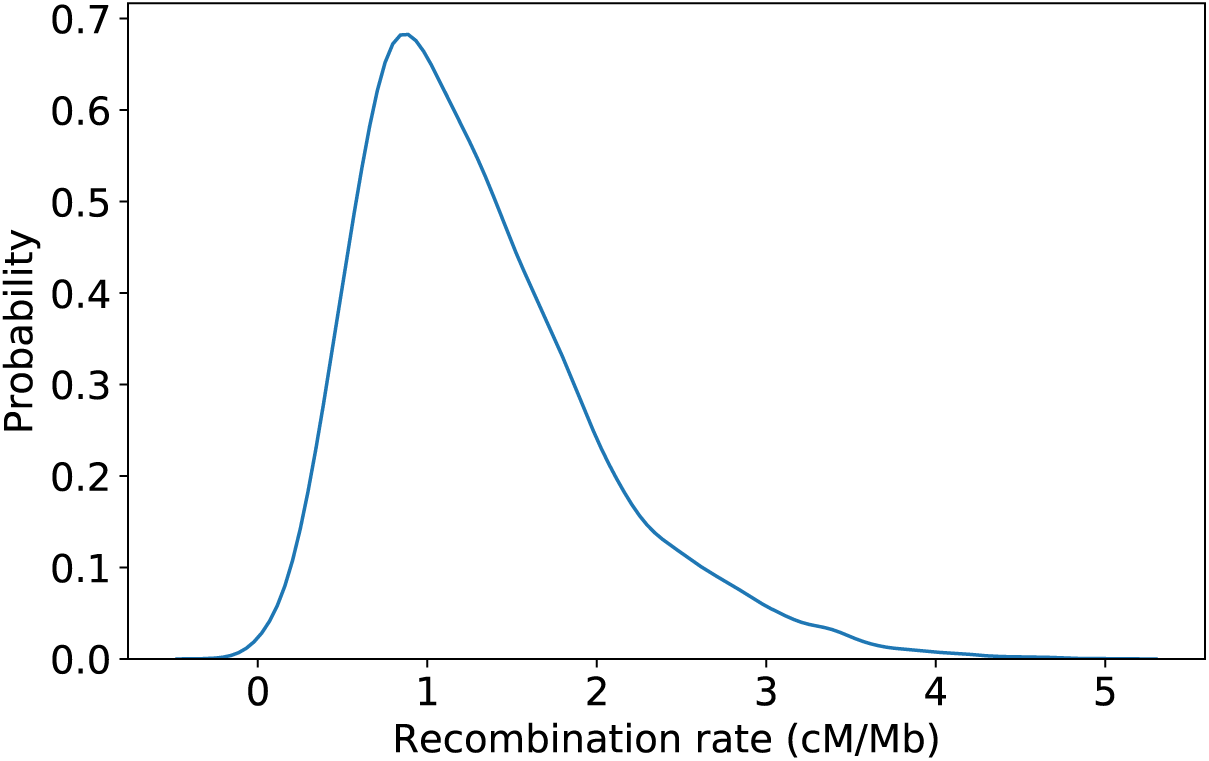
Human recombination rate distribution from which *ρ* is drawn. The recombination rate from the 1000 genomes project (Consortium et al, 2015) is averaged over 2Mb windows after masking centromeres and telomeres and fitted with a kernel Gaussian distribution.

### 1.3 Inferred population sizes by Attentive-SPIDNA on the HGDP dataset without interpolation

**Fig. S8:**
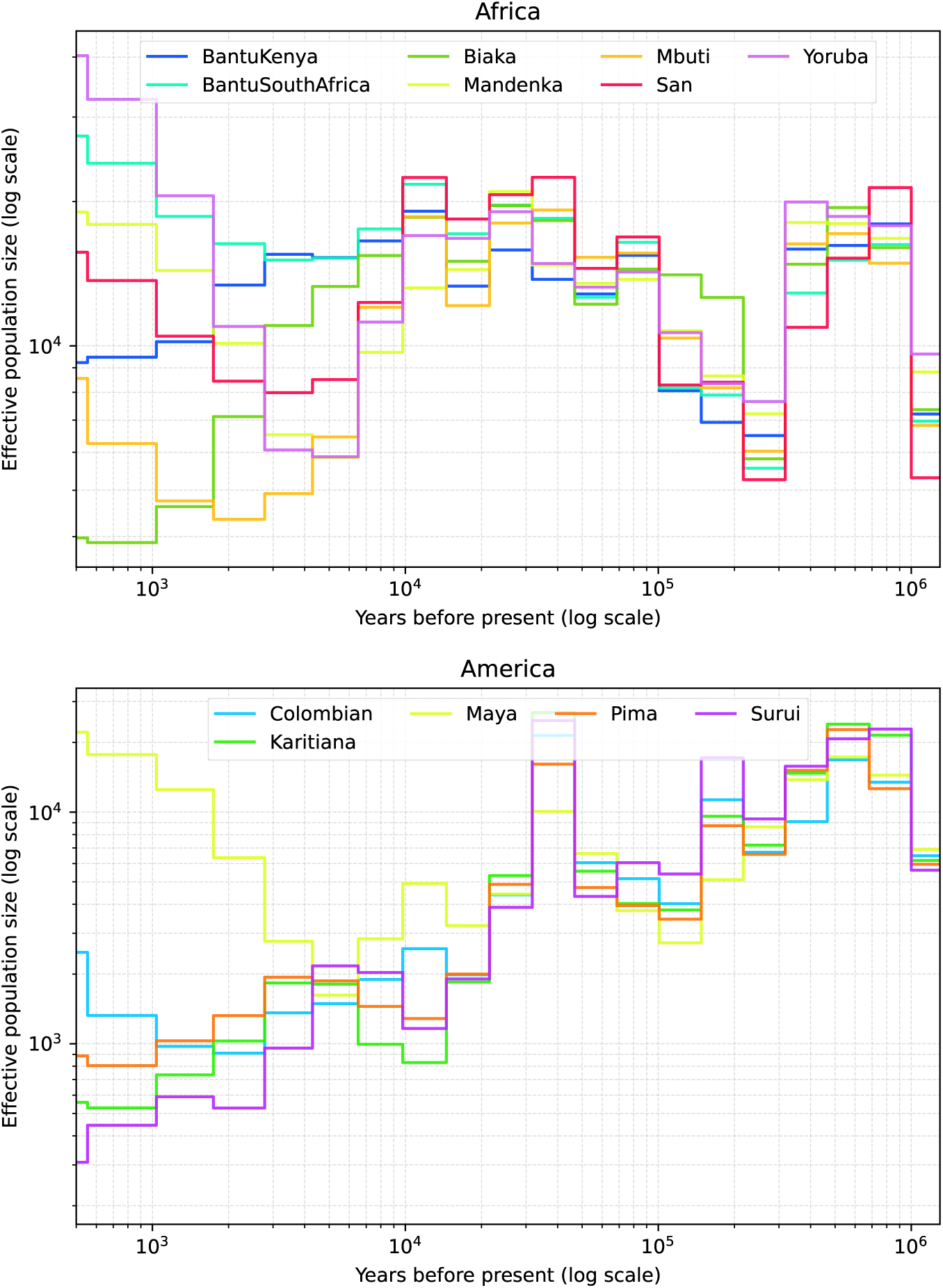
Effective population sizes inferred by Attentive-SPIDNA before interpolation on the HGDP dataset (1/4).

**Fig. S8:**
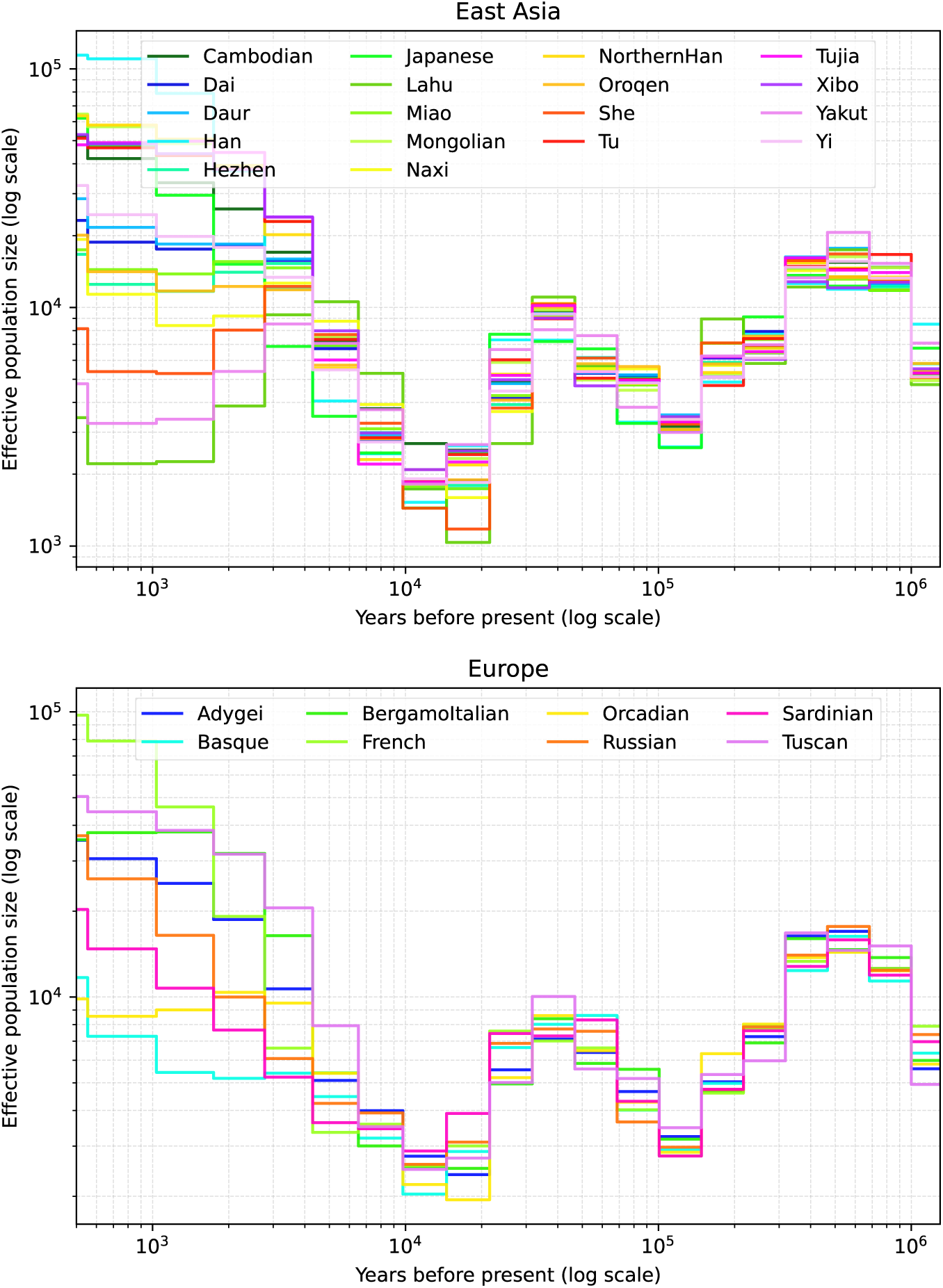
Effective population sizes inferred by Attentive-SPIDNA before interpolation on the HGDP dataset (2/4).

**Fig. S8:**
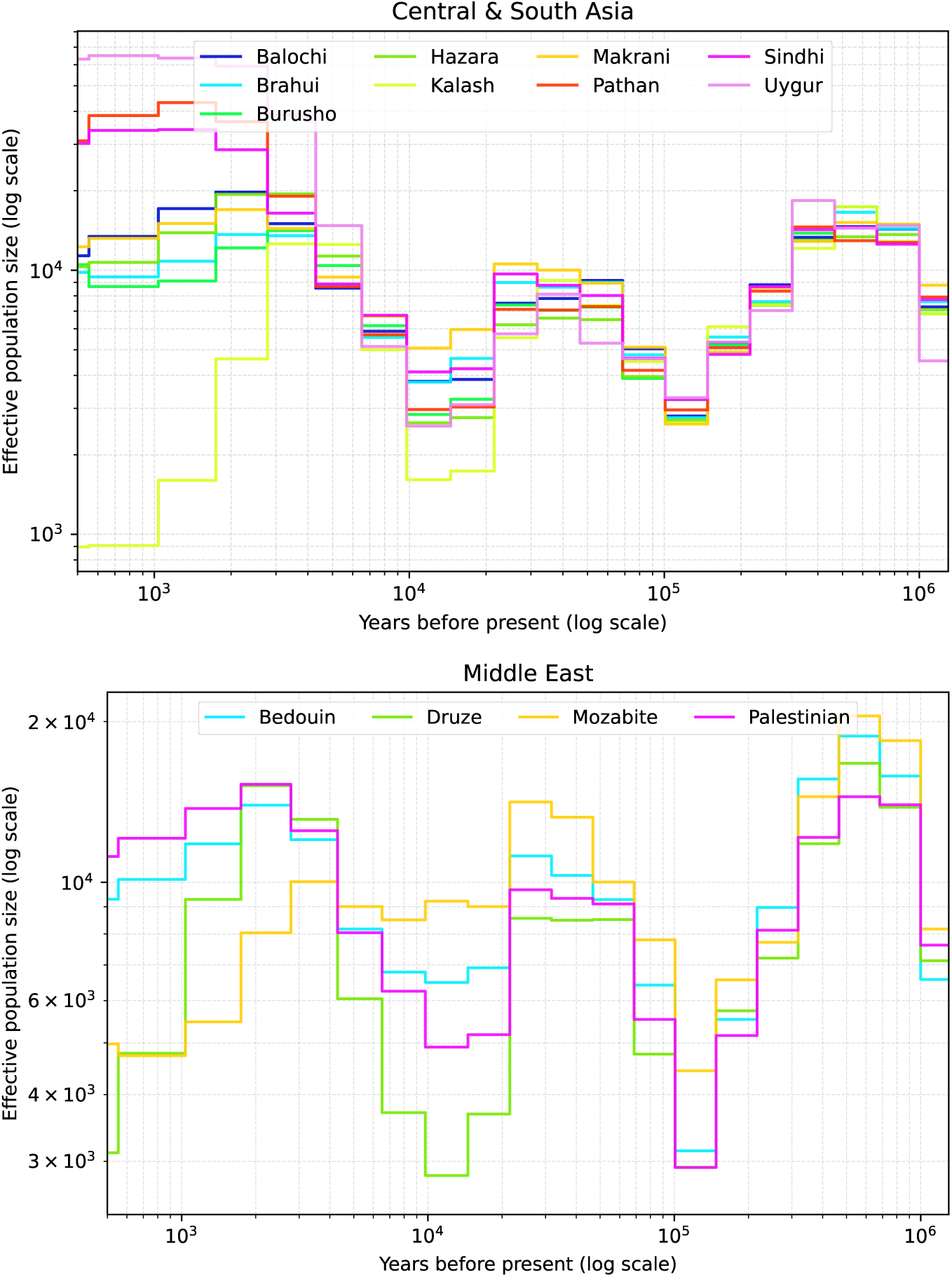
Effective population sizes inferred by Attentive-SPIDNA before interpolation on the HGDP dataset (3/4).

**Fig. S8:**
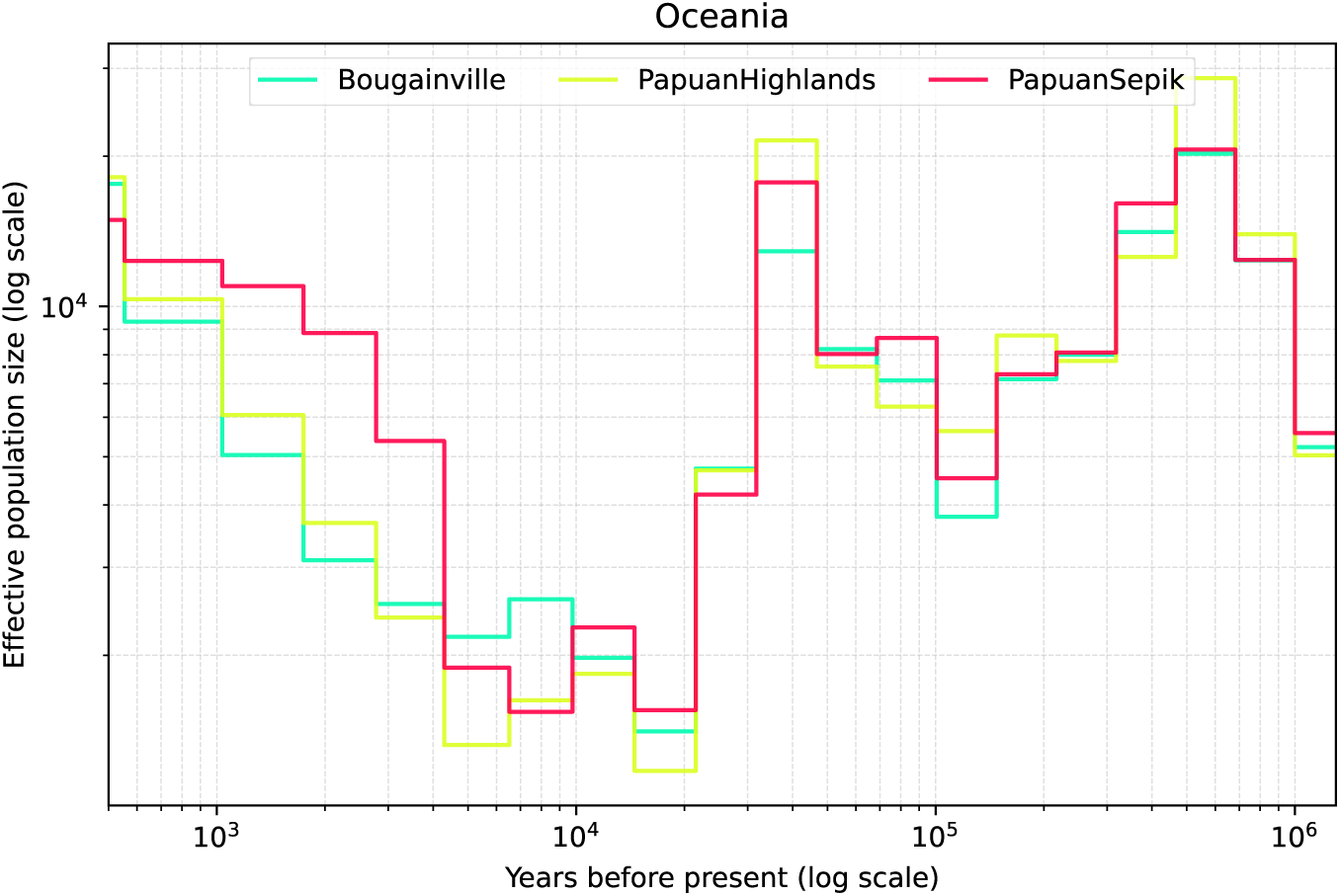
Effective population sizes inferred by Attentive-SPIDNA before interpolation on the HGDP dataset (4/4).

### 1.4 Robustness of Attentive-SPIDNA

#### 1.4.1 Robustness of Attentive-SPIDNA to the number of haplotype

**Fig. S9:**
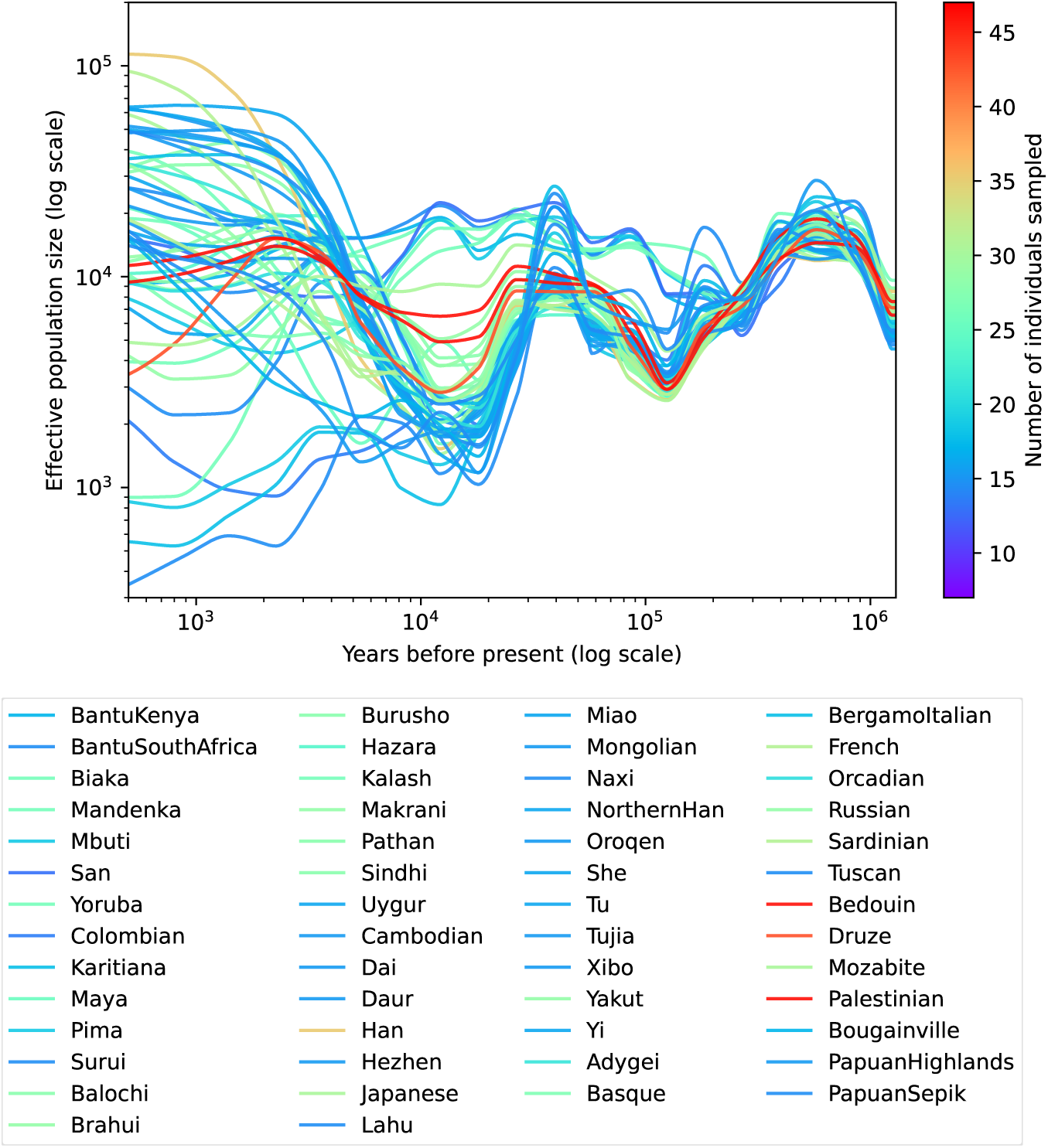
Effective population sizes inferred by Attentive-SPIDNA for the human populations from the HGDP dataset colored by the number of individuals sampled. Effective sizes are inferred by the Attentive-SPIDNA version with the lowest prediction error on the simulated dataset, i.e., with attention mechanism on scenario, batches with padding and the weight unfreezing mechanism. For each population, the 21 predicted values are interpolated using a piecewise cubic Hermite interpolator.

**Fig. S10:**
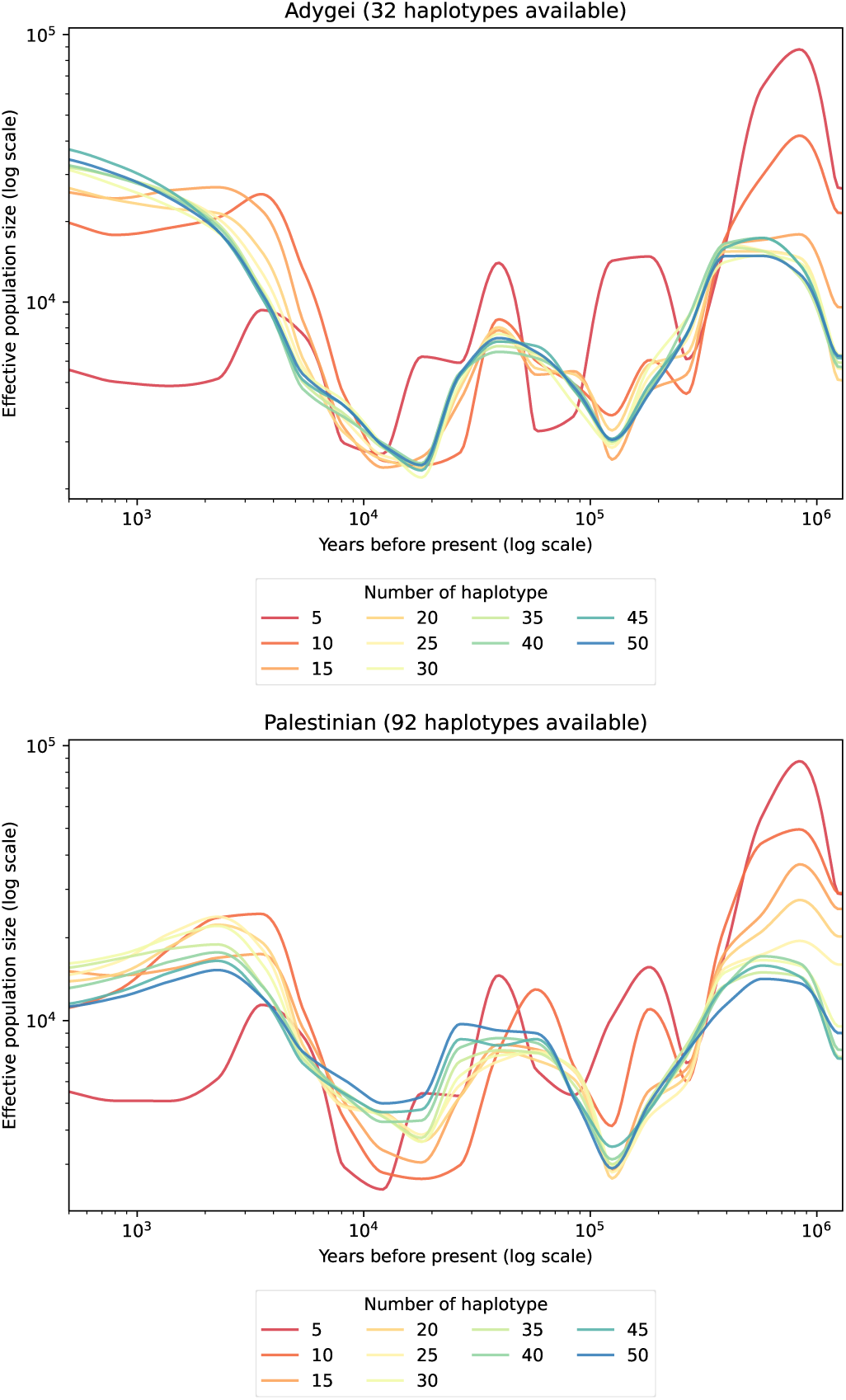
Attentive-SPIDNA effective population sizes inferred for Adygei and Palestinian populations with haplotype subsampling (1/2). Prediction of Attentive-SPIDNA for the Adygei and Palestinian populations when subsampling the number of haplotypes fed to A-SPIDNA from 50 to 5.

**Fig. S10:**
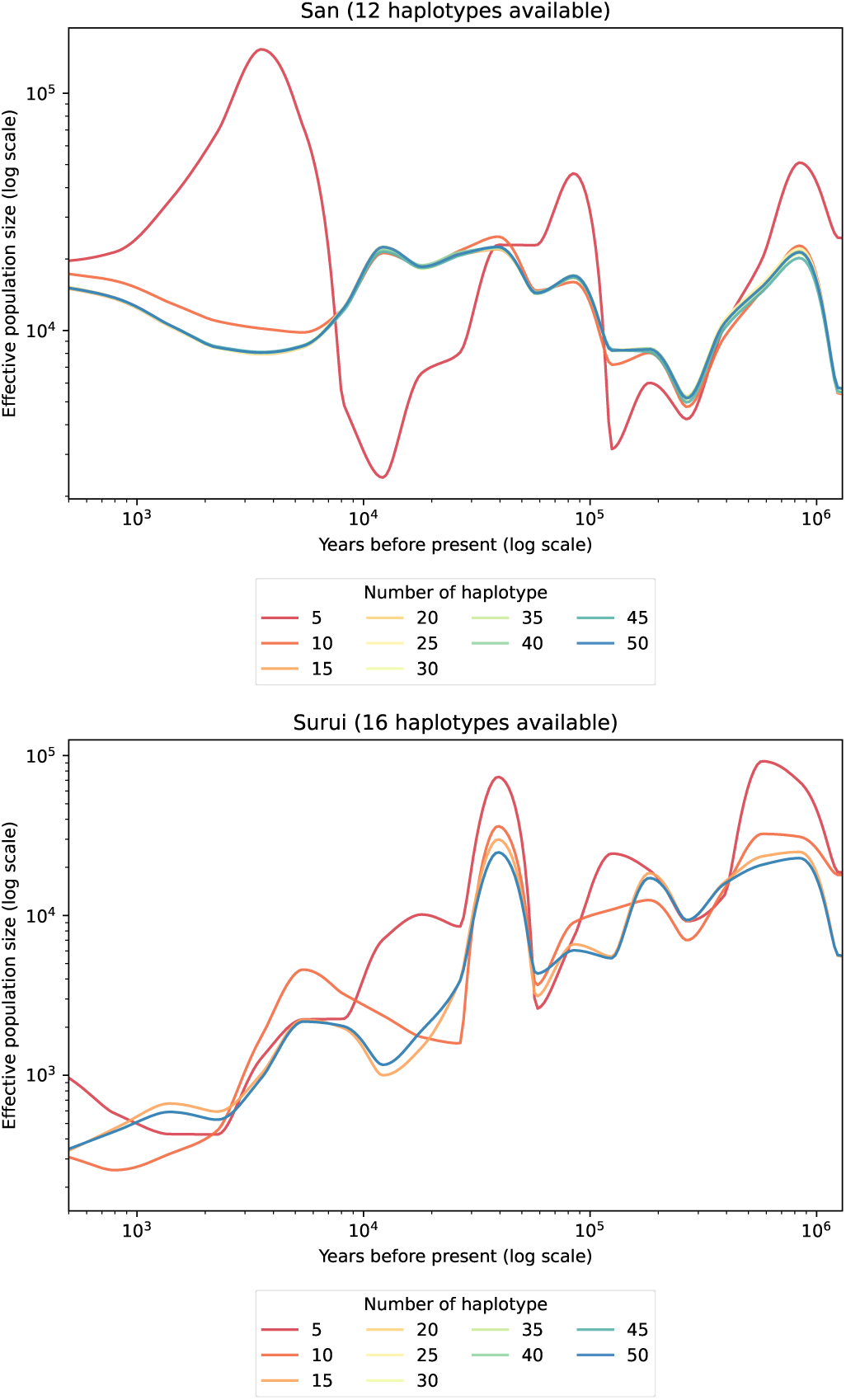
Attentive-SPIDNA effective population sizes inferred for San and Surui populations with haplotype subsampling (2/2). Prediction of Attentive-SPIDNA for the San and Surui populations when subsampling the number of haplotypes fed to A-SPIDNA from 50 to 5.

#### 1.4.2 Robustness of Attentive-SPIDNA to the batch size

**Fig. S11:**
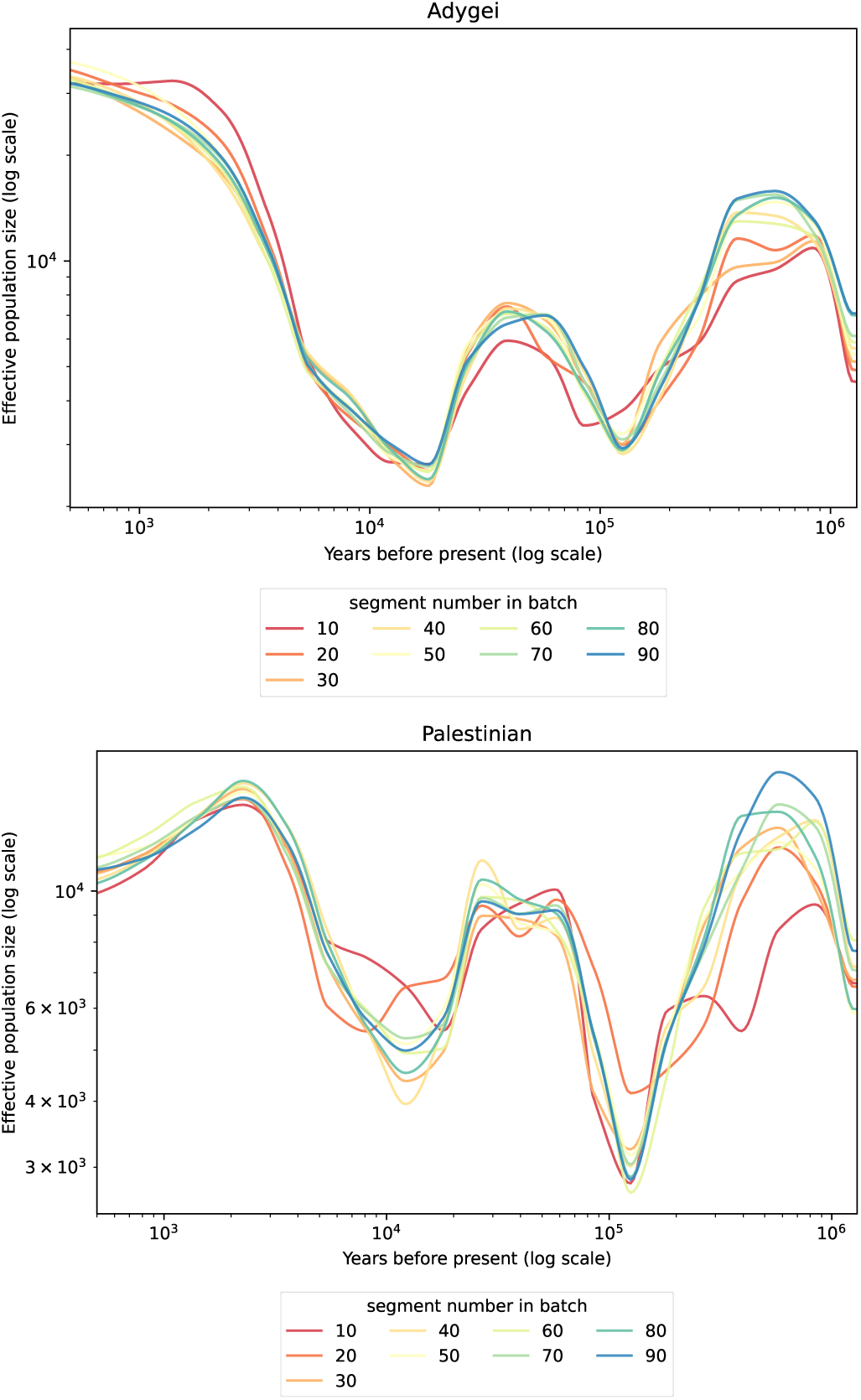
Attentive-SPIDNA effective population sizes inferred for Adygei and Palestinian populations with different batch sizes. Prediction of Attentive-SPIDNA of Adygei and Palestinian effective population sizes for batch sizes between 100 and 10 SNP matrices. Each line is the averaged prediction of 10 batches. Attentive-SPIDNA is trained with batches of 100 SNP matrices

### 1.5 Computational resources

Simulations have been performed on the genotoul bioinformatics platform with the following hardware:

- 68 nodes with 2 E5-2670 v2 Intel CPUs (2.50GHz, 20 threads) and 256GB of RAM
- 48 nodes with 2 E5-2683 v4 Intel CPUs (2.10GHz, 32 threads) and 256GB of RAM.

All summary statistics, trainings and predictions were computed on the TAU’s Titanic platform with the following hardware:

- 5 nodes with 4 GTX 1080 (12GB of VRAM) GPUs, 2 E5-2650 v4 Intel CPUs (2.20GHz, 24 threads) and 252GB of RAM
- 7 nodes with 4 RTX 2080 (12GB of VRAM) GPUs, 2 Silver 4108 Intel CPUs (1.80GHz, 18 threads) and 252GB of RAM
- 1 node with 4 Tesla P100 (16GB of VRAM) GPUs, 2 E5-2690 v4 Intel CPUs (2.60GHz, 28 threads) and 252GB of RAM
- 1 node with 2 RTX 2080 (8GB of VRAM) GPUs, 2 E5-2650 v4 Intel CPUs (2.20GHz, 24 threads) and 252GB of RAM

Both platforms use Slurm as job scheduling system. Batch sizes and deep learning architectures were all designed to fit on less than 12GB of VRAM during training. To train non-adaptive architectures, batches were split between 3 GPUs with at least 12GB of VRAM. Adaptive architectures were trained on one GPU as batch data of varying sizes could not be concatenated in the same tensor. The training of SPIDNA took at most 1h42 per epoch for non-adaptive version and 31h31 per epoch for adaptive version. The slow computation time of adaptive SPIDNA is mostly due to data from the same batch being inputted one by one in the network instead of concatenated in tensors.

